# Transcription factors that specify cell fate activate cell cycle regulator genes to determine cell numbers in ascidian larval tissues

**DOI:** 10.1101/2022.05.16.492211

**Authors:** Kenji Kobayashi, Miki Tokuoka, Hiroaki Sato, Manami Ariyoshi, Shiori Kawahara, Shigeki Fujiwara, Takeo Kishimoto, Yutaka Satou

## Abstract

In animal development, most cell types stop dividing before terminal differentiation; thus, cell cycle control is tightly linked to cell differentiation programs. Although cell cycle control in animal development has been studied extensively, such links are not well understood. In ascidian embryos, cell lineages do not vary among individuals, and rounds of the cell cycle are determined according to cell lineages. In the present study, we first showed that maternal factors promote approximately 10 rounds of cell division without zygotic gene expression. Notochord and muscle cells stop dividing after fewer than 10 rounds of cell division, and we show that a Cdk inhibitor (*Cdkn1.b*) is responsible. *Cdkn1.b* is also necessary for epidermal cells to stop dividing. In contrast, mesenchymal and endodermal cells divided more than 10 times, and *Myc*, which encodes a proto-oncogenic transcription factor, is responsible for maintaining cell cycle progression in these tissues. Expression of *Cdkn1.b* in notochord and muscle is controlled by the same developmental programs that specify the developmental fate of notochord and muscle. Likewise, expression of *Myc* in mesenchyme and endoderm was under control of the same developmental programs that specify the developmental fate of mesenchyme and endoderm. Because these transcription factors that regulate *Cdkn1.b* and *Myc* are essential factors for fate specification of these tissues, cell fate specification and cell cycle control are linked by those transcription factors. In other words, ectodermal, mesodermal, and endodermal tissues in ascidian embryos control the cell cycle through *Cdkn1.b* and *Myc*, which are under the control of transcription factors that specify cell fate.

## Introduction

In animal development, controlling the cell cycle is important for tissues and organs to be properly differentiated. In many tissues, terminal differentiation is coupled with withdrawal from the cell cycle (G0 arrest). For example, a myogenic bHLH transcription factor, Myog, induces cell cycle exit in differentiated myocytes in mammalian embryos (Liu et al., 2012). In nematode embryos, mutations in genes encoding cyclin-dependent kinase inhibitors induce hyperplasia in multiple somatic tissues, suggesting that these proteins normally stop cells from dividing (Fukuyama et al., 2003). Similarly, in fly embryos, *dacapo*, encoding a cyclin-dependent kinase inhibitor is necessary to stop epidermal cell proliferation (deNooij et al., 1996; Lane et al., 1996). In mice, Mycn is required to maintain proliferating neural cell progenitors, and a mutant of this gene resulted in precocious differentiation of neural cells (Knoepfler et al., 2002).

In embryos of an ascidian, *Ciona savignyi*, cell lineages are invariant among individuals, and numbers of cell divisions are strictly controlled (Conklin, 1905; Nishida, 1987). Notochord cells become postmitotic after nine rounds of cell cycle, and every larva contains exactly 40 notochord cells. Similarly, 36 muscle cells are differentiated in the larval tail, and the anterior 28 cells are derived from the posterior vegetal quadrant of an embryo. Among them, 16 cells become postmitotic after nine rounds of cell division, and 12 cells become postmitotic after eight rounds of cell division. Epidermal cells stop dividing after 11 rounds of cell division (Ogura et al., 2011; Pasini et al., 2006; Yamada and Nishida, 1999), although these cells resume cycling around the time when a larva obtains competence for metamorphosis (Nakayama et al., 2005). On the other hand, mesenchymal cells and endodermal cells, which produce adult tissues after metamorphosis, may continue to divide after hatching (Nakayama et al., 2005; Yamada and Nishida, 1999). In this way, these tissues in ascidian larvae regulate the cell cycle differently, which suggests that developmental programs for these tissues are involved in determining when cells stop dividing.

In the notochord lineage of embryos of *Halocynthia roretzi*, an ascidian belonging to a different order from that of *Ciona*, the number of cell divisions is controlled by a mechanism that is under control of *Brachyury* (or *T*) (Fujikawa et al., 2011), a key transcription factor for notochord differentiation (Chiba et al., 2009; Satou et al., 2001a; Yasuo and Satoh, 1994). Similarly, the number of cell divisions in muscle cells is controlled by *Tbx6-r* (Kuwajima et al., 2014), a key transcription factor for muscle differentiation (Yagi et al., 2005; Yu et al., 2019). Because a gene encoding a Cdk inhibitor (*CKI-b* or *Cdkn1.b*) is expressed in notochord and muscle cells, and because expression in the notochord is under control of *Brachyury*, this gene may be involved in regulation of cell cycle in these tissues. Indeed, injection of *Cdkn1.b* mRNA into fertilized eggs halts the cell cycle around the 8- or 16-cell stage (Kuwajima et al., 2014), although its function in differentiation of notochord and muscle has not been evaluated.

On the other hand, mesenchymal and endodermal cells likely continue to divide even after hatching (Nakayama et al., 2005), and produce a large number of small cells; however, it is not known how these tissues maintain cell cycle progression. It is also uncertain whether *Twist-r.a/b* and *Lhx3/4*, which encode key transcription factor genes for fate specification of mesenchyme and endoderm (Imai et al., 2003; Satou et al., 2001b), control cell numbers in these tissues. In the present study, we addressed these issues and found that *Cdkn1.b* and *Myc* play key roles in cell cycle control in ascidian embryos.

## Results

### *Ciona* fertilized eggs divide approximately ten times without zygotic transcription

To test how many rounds of cell division occur without zygotic transcription, we injected α-amanitin, which inhibits RNA polymerase II, into eggs of ascidians, *C. savignyi*. When uninjected control eggs developed to larvae, we fixed four α-amanitin-injected embryos. The mean number of nuclei in these embryos was 1306.5 (Figure 1A, B). This may suggest that each cell divided 10 or 11 times from the 1-cell stage, if we assume that there are no cells that divide much more frequently than others. We also counted nuclei of three embryos that developed in sea water containing actinomycin D, another inhibitor of transcription. The average number was 584.7 (Figure 1C, D).

**Figure 1.**
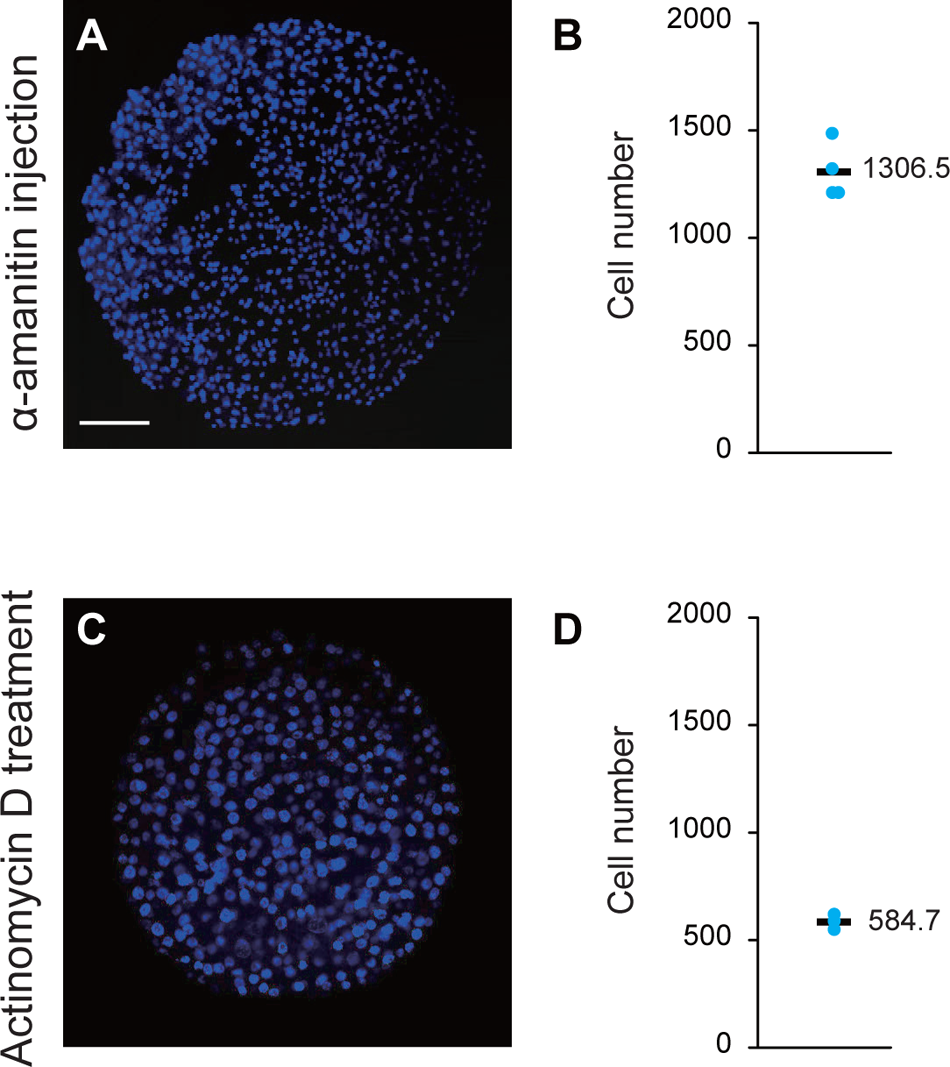
Maternal factors can promote 9 to 11 rounds of cell division without zygotic gene expression. (A) An embryo developed from an egg injected with α-amanitin. This embryo was fixed when uninjected control larvae hatched. Nuclei were stained with DAPI. The scale bar represents 50 μm. (B) Four dots indicate numbers of nuclei of four embryos developed from eggs injected with α-amanitin. A bar indicates the average. (C) An embryo developed in sea water containing actinomycin C. Nuclei were stained with DAPI. (D) Three dots indicate numbers of nuclei of three embryos incubated in sea water containing Actinomycin D. A bar indicates the average.

This suggests that each cell divided 9 or 10 times from the 1-cell stage. Although it was not clear why these two drugs yielded slightly different results, these drug treatment experiments showed that cells divided 9–11 times in embryos in which transcription was inhibited. In other words, maternal factors can induce nine to eleven rounds of cell division. Because cell cycle rounds of most ascidian larval cells are strictly controlled according to their cell lineages, this result suggests that zygotically expressed genes control numbers of cell divisions.

### Zygotic gene expression is necessary for notochord, muscle, and epidermal cells to stop dividing

Notochord cells stop dividing after nine rounds of cell division, and epidermal cells stop after 11 rounds of cell division (Nishida, 1987; Ogura et al., 2011; Pasini et al., 2006; Yamada and Nishida, 1999). The primary lineage of muscle cells (the anterior 28 muscle cells) stop dividing after 8 or 9 rounds of cell division depending on their lineages (Nishida, 1987). To examine whether zygotic gene expression is involved in regulation of cell cycle counts, we isolated cells of presumptive notochord and muscle at the 64-cell stage, and presumptive epidermal cells at the 32-cell stage using a fine glass needle.

First, we isolated one presumptive notochord cell (A7.3/A7.7) from each of 11 normal 64-cell embryos. When isolated cells were incubated in normal sea water until uninjected control embryos hatched, they divided three times to become eight cells in most cases (Figure 2A). Because each of these cells produces eight notochord cells in normal embryos, this observation indicated that the number of cell cycle rounds in these isolated cells was regulated autonomously, as previously indicated in *Halocynthia* embryos (Fujikawa et al., 2011). When we isolated presumptive notochord cells from embryos injected with α-amanitin, each of the isolated cells became 15.2 cells on average. Similarly, when isolated cells were incubated in sea water containing actinomycin D, each of these cells became 13.4 cells on average (Figure 2A). Thus, inhibitors of transcription increased rounds of the cell cycle of isolated presumptive notochord cells, indicating that zygotic gene expression was required to stop cell division in the notochord lineage.

**Figure 2.**
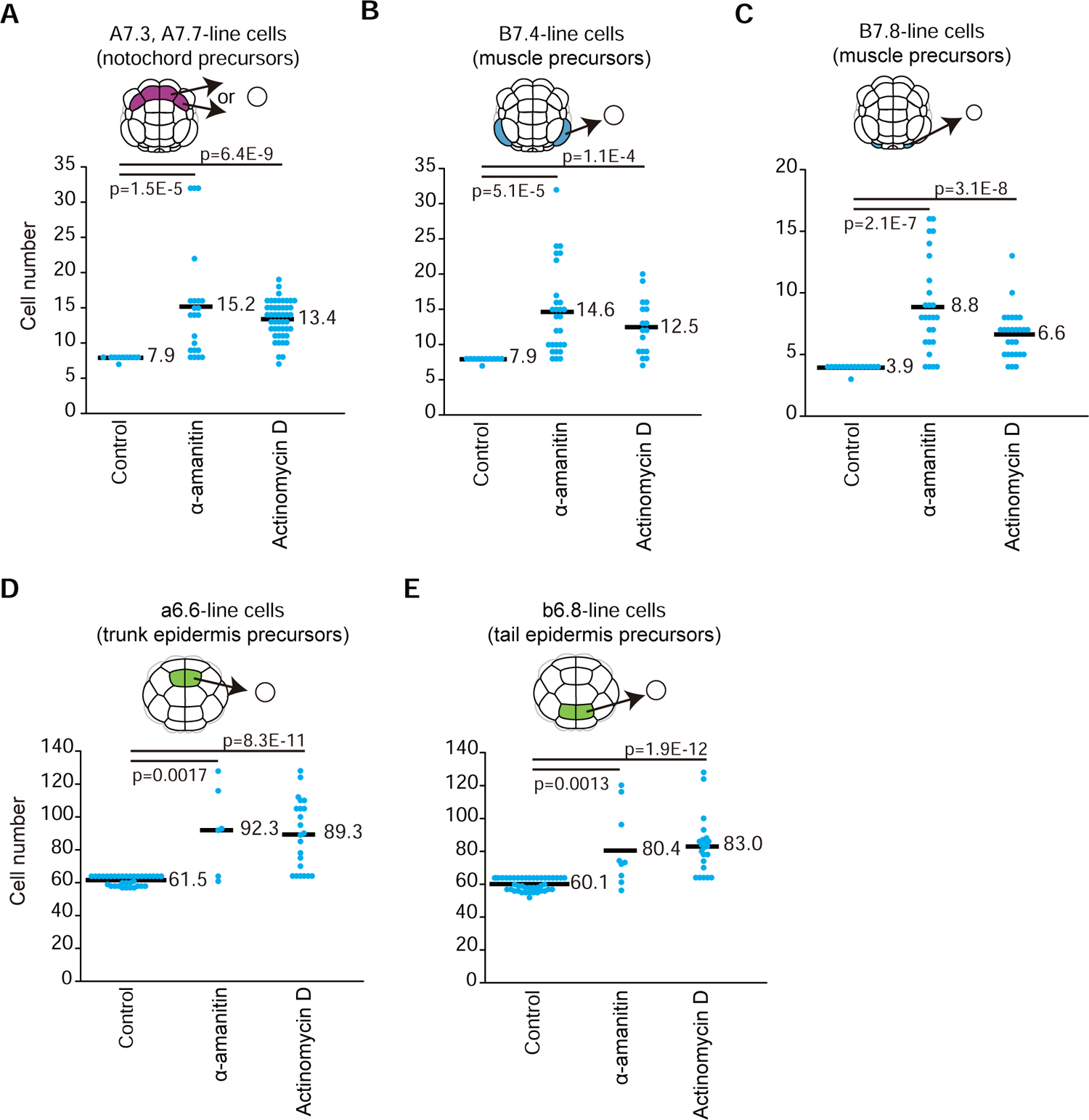
The number of cells produced from an isolated presumptive notochord, muscle, or epidermal cell. (A-C) At the 64-cell stage, one presumptive notochord cell (A7.3 or A7.7; A) or presumptive muscle cell (B7.4 in B or B7.8 in C) was isolated from each of normal embryos, embryos injected with α-amanitin, and embryos incubated in sea water containing actinomycin D. Partial embryos were incubated until uninjected control embryos hatched. Partial embryos obtained from actinomycin D-treated embryos were incubated in sea water containing actinomycin D until fixation. Numbers of cells in resulting partial embryos were counted. Embryos were injected with α-amanitin or incubated in sea water containing actinomycin D. Differences in cell number were examined with the two-sided Wilcoxon rank sum test, and p-values are indicated. Each dot indicates the number of labelled cells in a single embryo, and averages are shown by black bars. (D, E) Presumptive epidermal cells (a6.6 in D; b6.8 in E) were similarly isolated at the 32-cell stage, and counted.

Similarly, we isolated two different lineages of presumptive muscle cells from normal 64-cell embryos. First, when B7.4 cells were isolated and incubated in normal sea water, isolated cells divided three times to become 7.9 cells on average (Figure 2B). This number was close to the expected number (8=2^3^), because B7.4 cells divide three times before stopping division in normal embryos (Nishida, 1987). Therefore, as in notochord and muscle cells in *Halocynthia* embryos (Fujikawa et al., 2011), this experiment showed that the number of cell cycle rounds of isolated blastomeres was autonomously determined. On the other hand, when we isolated this lineage of cells from embryos injected with α-amanitin, each isolated cell became 14.6 cells. Similarly, when isolated cells were incubated in sea water containing actinomycin D, each of them became 12.5 cells on average (Figure 2B). Thus, zygotic gene expression was also required to stop cell division in the muscle lineage.

Similarly, a previous study reported that B7.8 presumptive muscle cells divide two additional times before stopping division in normal embryos (Nishida, 1987), and we found that isolated B7.8 cells usually divide twice in normal sea water (Figure 2C). However, B7.8 cells from α-amanitin-injected embryos became 8.8 cells on average, and isolated B7.8 cells incubated in sea water containing actinomycin D became 6.6 cells on average (Figure 2C). Therefore, zygotic gene expression is also necessary to stop the cell cycle in this muscle lineage.

Finally, we isolated presumptive epidermal cells (a6.6 and b6.8) from 32-cell embryos. These cells divide six more times (11 times in total from 1-cell embryos) in normal embryos. When isolated a6.6 and b6.8 cells were incubated in normal sea water, they became 61.5 and 60.1 cells on average. These numbers were close to the expected value (64=2^6^) (Figure 2D, E). On the other hand, a6.6 and b6.8 cells isolated from embryos injected with α-amanitin became 92.3 and 80.4 cells, and a6.6 and b6.8 cells incubated in sea water containing actinomycin D became 89.3 and 83.0 cells (Figure 2D, E). Therefore, cell cycle rounds of epidermal cells were also under control of zygotic gene expression.

### A Cdk inhibitor controls the number of cell divisions of notochord, muscle, and epidermal cells

Involvement of a Cdk inhibitor in regulation of cell division of notochord and muscle cells has been indicated in another ascidian (Kuwajima et al., 2014). We identified an ortholog (*Cdkn1.b*) and examined its expression pattern by *in situ* hybridization (Figure 3). No clear signal for zygotic expression was observed before the gastrula stage, and the first evident *in situ* hybridization signal was detected in primary muscle lineage cells at the middle gastrula stage. Soon after, presumptive notochord cells expressed *Cdkn1.b*. Signals were also detected in a few neural cells at the early neurula stage, and many neural cells expressed *Cdkn1.b* at the middle and late neurula stages. At the late neurula stage, signals in epidermal cells also became evident transiently. At the early and middle tailbud stages, signals in notochord, muscle, and neural cells were evident, while signals in epidermal cells became undetectable.

**Figure 3.**
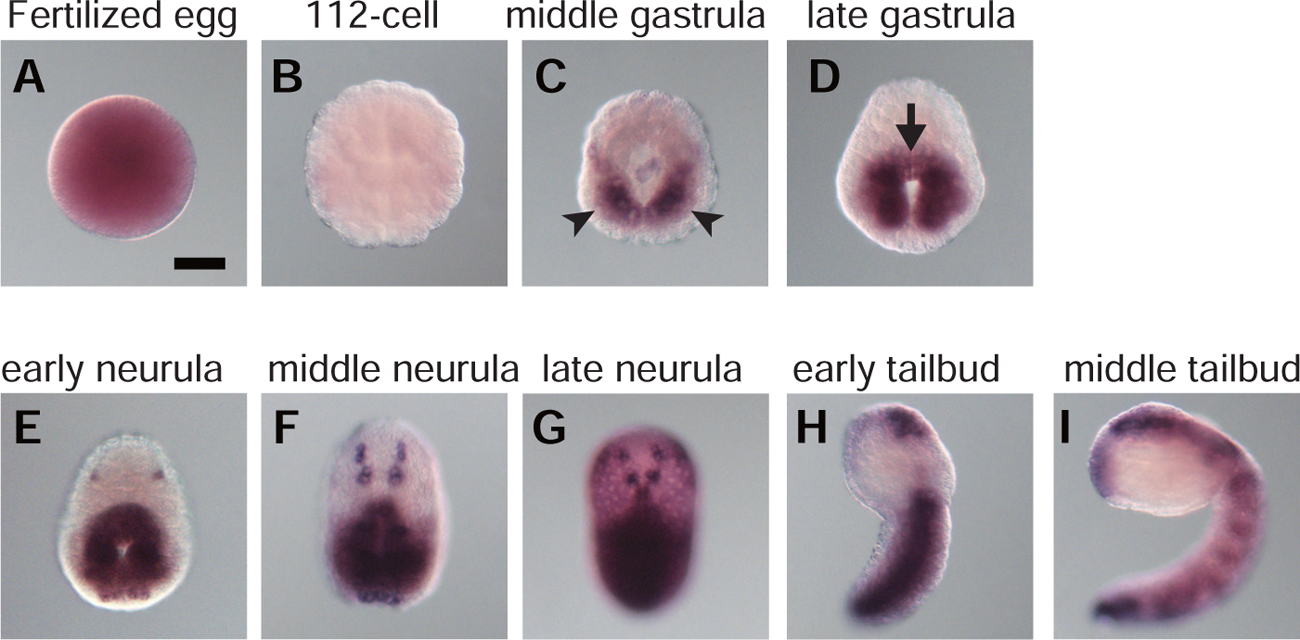
Expression pattern of *Cdkn1.b* in *C. savignyi* embryos. Photographs of whole mount *in situ* hybridization of (A) a fertilized egg and embryos at the (B) 112-cell, (C) middle gastrula, (D) late gastrula, (E) early neurula, (F) middle neurula, (G) late neurula, (H) early tailbud and (I) middle tailbud stages. Expression in the muscle lineage was first evident at the middle gastrula stage (black arrowheads). Expression in the notochord lineage was first evident at the late gastrula stage (black arrow). Expression in epidermal cells was transiently evident at the late neurula stage. Scale bar, 50 μm.

Next, we injected a morpholino antisense-oligonucleotide (MO) into unfertilized eggs, which was designed to inhibit translation of *Cdkn1.b* mRNA. After insemination, injected eggs were incubated until the 64-cell stage. Then, to count how many times cells divided, we isolated presumptive notochord cells (A7.3 or A7.7), presumptive muscle cells (B7.4 and B7.8), and presumptive epidermal cells (a6.6 and b6.8), and incubated them until uninjected control embryos hatched. As expected, isolated cells from *Cdkn1.b*-MO-injected embryos produced a larger number of cells than isolated cells from embryos injected with a control MO against *Escherichia coli lacZ*, and the differences were statistically significant in all cases (Figure 4A-E). These numbers were scarcely changed when we incubated for an additional 6 h. These results strongly supported the previously presented hypothesis that *Cdkn1.b* is responsible for stopping cell division in notochord and muscle (Kuwajima et al., 2014). In addition, the present study showed that *Cdkn1.b* is also transiently expressed in epidermal cells in order for epidermal cells to stop dividing.

**Figure 4.**
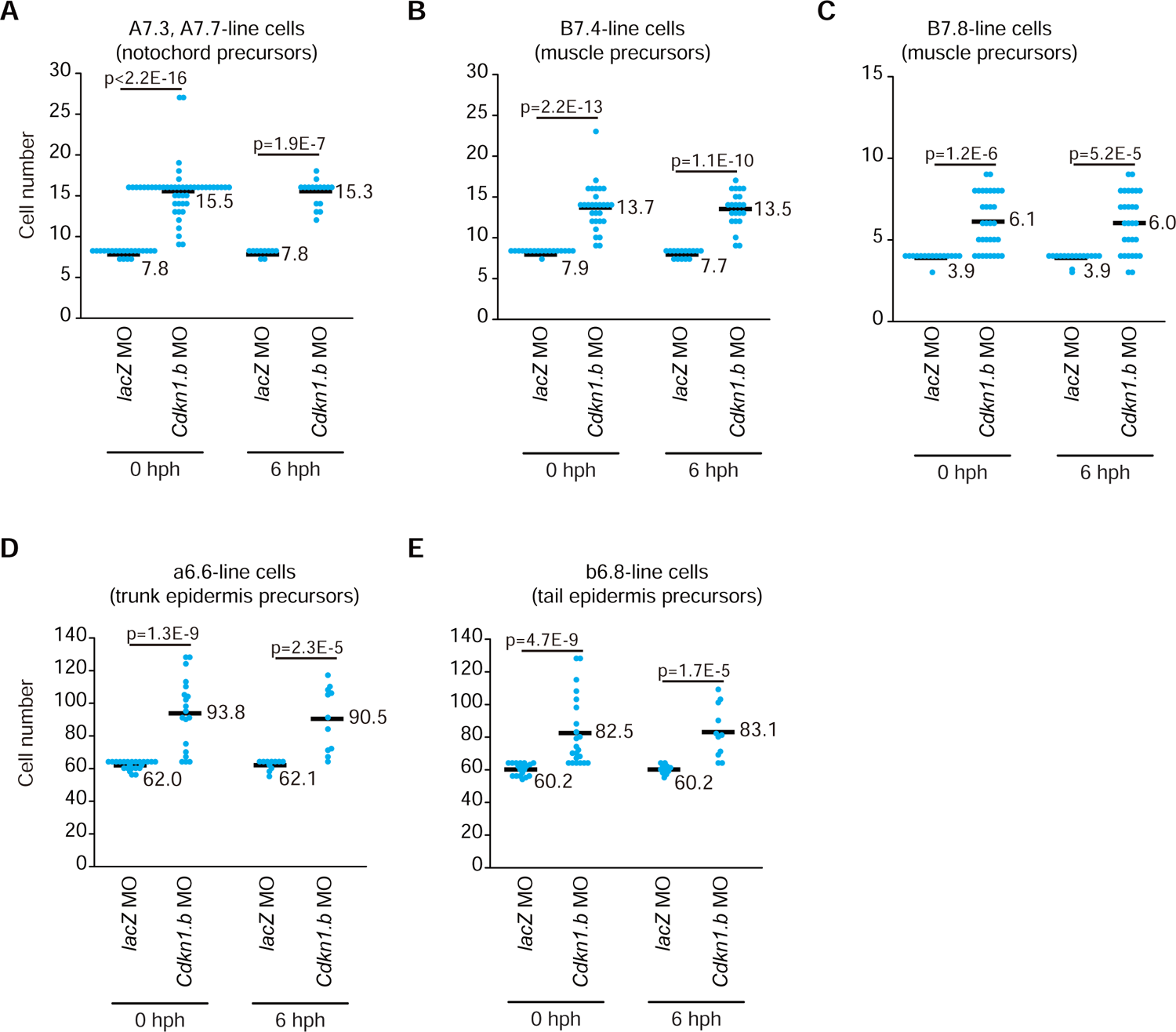
Presumptive notochord, muscle, or epidermal cells isolated from *Cdkn1.b* morphant embryos produced larger numbers of cells than those isolated from control embryos. The control *lacZ* or *Cdkn1.b* MO was injected into eggs. At the 32-or 64-cell stage, a presumptive notochord (A7.3 or A7.7; A), muscle (B7.4, B; B7.8, C), or epidermal cell (a6.6, D; b6.8, E) was isolated and incubated until uninjected sibling embryos hatched (0 hph) or 6 h later (6 hph). Embryos were fixed and cell numbers were counted. Differences in cell number were examined with the two-sided Wilcoxon rank sum test, and p-values are indicated. Each dot indicates the number of labelled cells in a single embryo, and averages are shown by black bars.

To confirm the specificity of the MO, we designed another MO against *Cdkn1.b*. We isolated presumptive notochord (A7.3/A7.7), and muscle (B7.4) from embryos injected with the second *Cdkn1.b* MO, and found that injection of this MO similarly produced a larger number of cells than the control MO (Figure S1).

### *Cdkn1.b* is under the control of *Brachyury*, *Zic-r.b*, and *Zic-r.a* in notochord and muscle cells

Because *Brachyury* directly or indirectly controls many notochord-specific genes (Takahashi et al., 1999), we examined whether expression of *Cdkn1.b* is under control of *Brachyury* in notochord. *Cdkn1.b* expression was indeed lost in presumptive notochord cells of embryos injected with a *Brachyury* MO, while it was unchanged in embryos injected with the control MO (Figure 5A, B). This observation is consistent with what was shown in *Halocynthia* embryos (Kuwajima et al., 2014). Next, we injected an MO against *Zic-r.b* (or *ZicL*). Because knockdown of *Zic-r.b* abolishes *Brachyury* expression (Imai et al., 2002), it was expected that knockdown of *Zic-r.b* would also suppress *Cdkn1.b*. Indeed, *Cdkn1.b* expression was lost in these embryos (Figure 5C).

**Figure 5.**
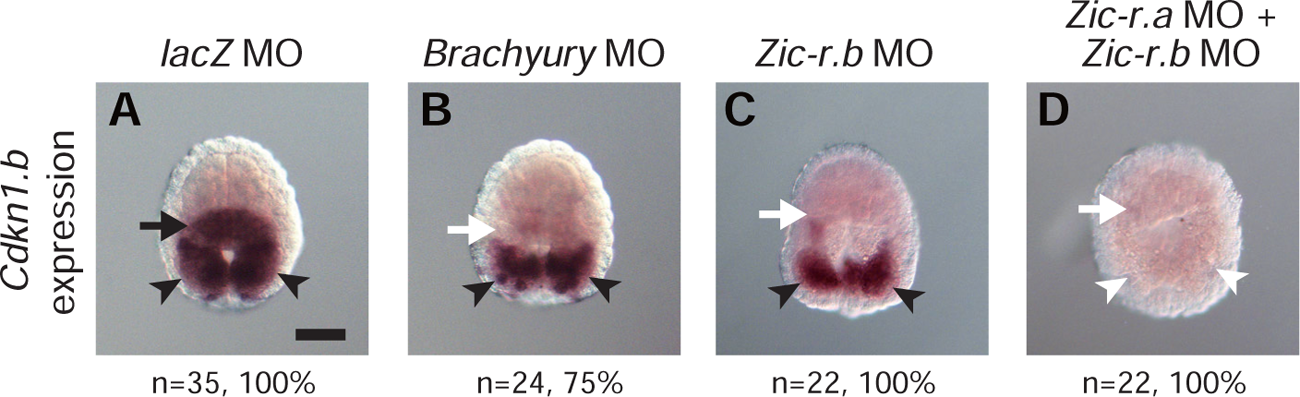
*Cdkn1.b* is controlled by *Brachyury*, *Zic-r.b*, and *Zic-r.a*. *In situ* hybridization of *Cdkn1.b* at the late gastrula stage in embryos injected with the (A) control *lacZ* MO, (B) *Brachyury* MO, and (C) *Zic-r.b* MO. The embryo in (D) was developed from an egg injected with two MOs for *Zic-r.a* and *Zic-r.b* MO simultaneously. Presumptive notochord and muscle cells are indicated by arrows and arrowheads, respectively. Black arrows and arrowheads indicate expression of *Cdkn1.b*, and white arrows and arrowheads indicate that *Cdkn1.b* expression was lost or decreased. Numbers of embryos examined and the proportion of embryos that each photograph represents are shown below the panels. Scale bar, 50 μm.

*Zic-r.b* is also involved in specification of muscle fate by activating *Tbx6-r.b* and *Mrf* in cooperation with a paralogous gene, *Zic-r.a* (or *Macho-1*) (Nishida and Sawada, 2001; Yu et al., 2019). While knockdown of *Zic-r.a* or *Zic-r.b* does not completely abolish expression of muscle-specific genes, simultaneous knockdown of *Zic-r.a* and *Zic-r.b* abolishes it completely (Imai et al., 2002; Satou et al., 2002). Similarly, while knockdown of *Zic-r.b* alone did not affect *Cdkn1.b* expression in muscle cells (Figure 5C), double knockdown of *Zic-r.a* and *Zic-r.b* abolished *Cdkn1.b* expression in muscle cells (Figure 5D). Thus, *Cdkn1.b* is controlled by the same developmental programs that direct fate specification in notochord and muscle.

Finally, to demonstrate the importance of cell cycle control in proper formation of notochord, we injected the *Cdkn1.b* MO into a pair of anterior vegetal blastomeres (A4.1 cell pair) of 8-cell embryos, because this cell pair produces 32 notochord cells, but does not produce primary lineage muscle cells or epidermal cells. Resulting larvae had short and thick tails, compared to normal larvae (Figure 6A, B). In early tailbud embryos injected with the control MO, 40 notochord cells were intercalated and aligned in a line (Figure 6C). On the other hand, in *Cdkn1.b*-MO-injected embryos, notochord cells were not aligned, although the number of notochord cells was the same as that in normal embryos at this stage (Figure 6D). Because major morphogenetic events of notochord formation start after notochord cells become postmitotic (Jiang and Smith, 2007), it seemed likely that inhibition of *Cdkn1.b* would also prevent notochord morphogenesis. Meanwhile, we examined expression of two marker genes, *Talin* (Satou et al., 2001c; Yamada et al., 2003) and *Tropomyosin-like 1* (Di Gregorio and Levine, 1999), by *in situ* hybridization (Figure 6E-H). As shown in Figure 6F, H, these markers were expressed in *Cdkn1.b* morphant embryos, indicating that knockdown of *Cdkn1.b* did not affect expression of *Talin* or *Tropomyosin-like 1* in notochord cells. Similarly, we confirmed that muscle markers and epidermal markers were expressed in *Cdkn1.b* morphant embryos (Figure S2).

**Figure 6.**
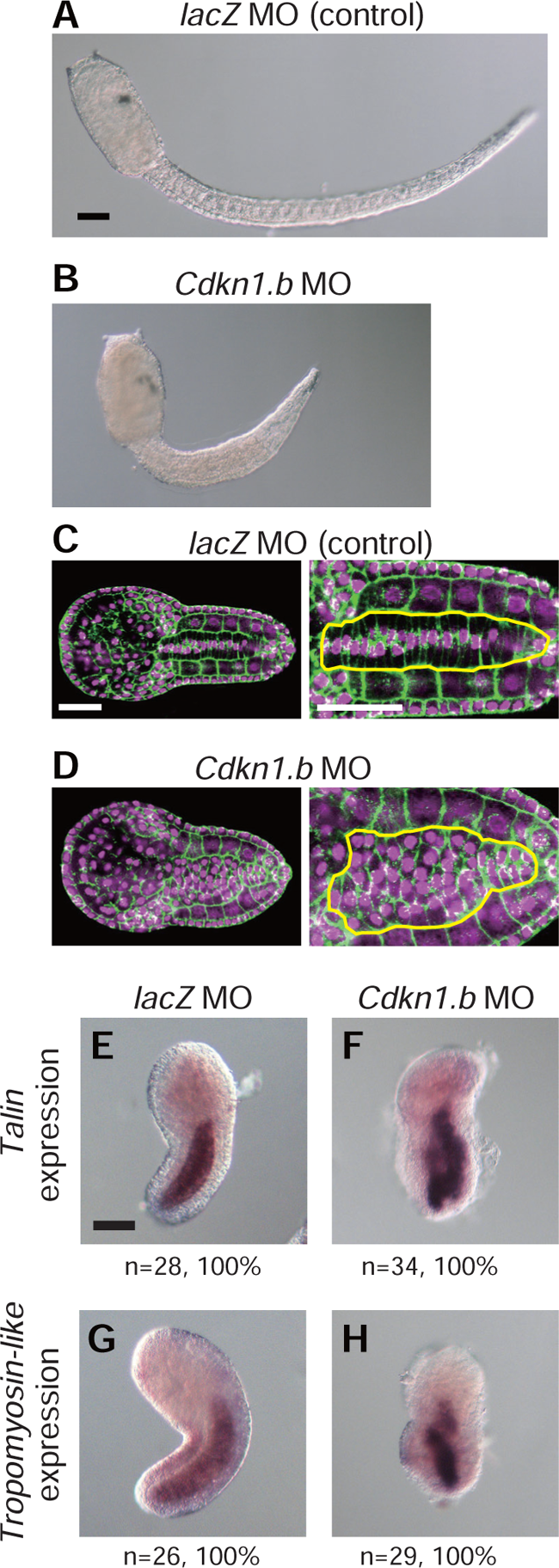
Notochord cells in embryos injected with the *Cdkn1.b* MO. (A, B) We injected the (A) control *lacZ* MO and (B) *Cdkn1.b* MO into a pair of anterior vegetal blastomeres (A4.1 cell pair) of 8-cell embryos, and a resultant larva is shown. (C, D) We similarly injected the control *lacZ* MO (C) or *Cdkn1.b* MO (D) into the A4.1 pair. Embryos were fixed at the early tailbud stage, and stained with phalloidin (green). The tail region is magnified in the right panels. In contrast to control embryos, notochord cells are not properly aligned in a line in *Cdkn1.b* MO-injected embryos. Nuclei were stained with DAPI (magenta). Notochord is enclosed by yellow lines. (E–H) Expression of *Talin* (E, F) and *Tropomyosin-like 1* (G, H) in early tailbud embryos injected with the control *lacZ* MO (E, G) or *Cdkn1.b* MO (F, H). Although morphology was severely disrupted, these notochord marker genes were expressed. Scale bars, 50 μm.

### *Myc* is required for mesenchymal and endodermal cells to continue to divide

A previous study showed that mesenchymal and endodermal cells in *Ciona robusta* (or *Ciona intestinalis* type A) continue to divide after hatching (Nakayama et al., 2005). We confirmed that these cells continue to divide after hatching in *C. savignyi* by labelling one of three presumptive mesenchyme cells (B7.7, B8.5, and A7.6) and five presumptive endoderm cells (B7.1, B7.2, A7.1, A7.2, and A7.5) with DiI. Numbers of labelled cells in larvae 6 h post-hatch (hph) were greater than those at hatching (Figure S3A, B).

For further confirmation, we transferred normal larvae into sea water containing BrdU for 30 min before fixation. At 0 and 6 hph, incorporation of BrdU in mesenchymal and endodermal cells was confirmed with a specific antibody (Figure S3C).

A previous study showed that a proto-oncogene, *Myc*, is expressed in mesenchyme and endoderm in *C. robusta* (or *C. intestinalis* type A) (Imai et al., 2004), and we confirmed that *Myc* was expressed in the same way in *C. savignyi* (Figure S4). On the basis of this expression pattern, we tested the possibility that *Myc* is involved in cell cycle regulation of mesenchymal and endodermal cells. For this purpose, we injected an MO against *Myc* into eggs, and labelled a presumptive mesenchyme or endoderm cell with DiI between the 68-cell and 112-cell stages. When the control *lacZ* MO was injected and B7.7 (presumptive mesenchymal cell) was labeled with DiI, 107.4 and 117.4 labelled cells were detected at 0 hph and 6 hph, respectively (Figure 7A). On the other hand, when the *Myc* MO was injected, 27.1 and 28.4 labelled cells were observed at 0 hph and 6 hph (Figure 7A). This suggested that control larvae divided approximately twice more than *Myc-* morphant embryos. We obtained similar results in the two remaining mesenchymal lineages (B8.5 and A7.6-lineages; Figure S5A). Therefore, the above observation indicated that *Myc* is necessary for mesenchyme cells to continue to divide.

**Figure 7.**
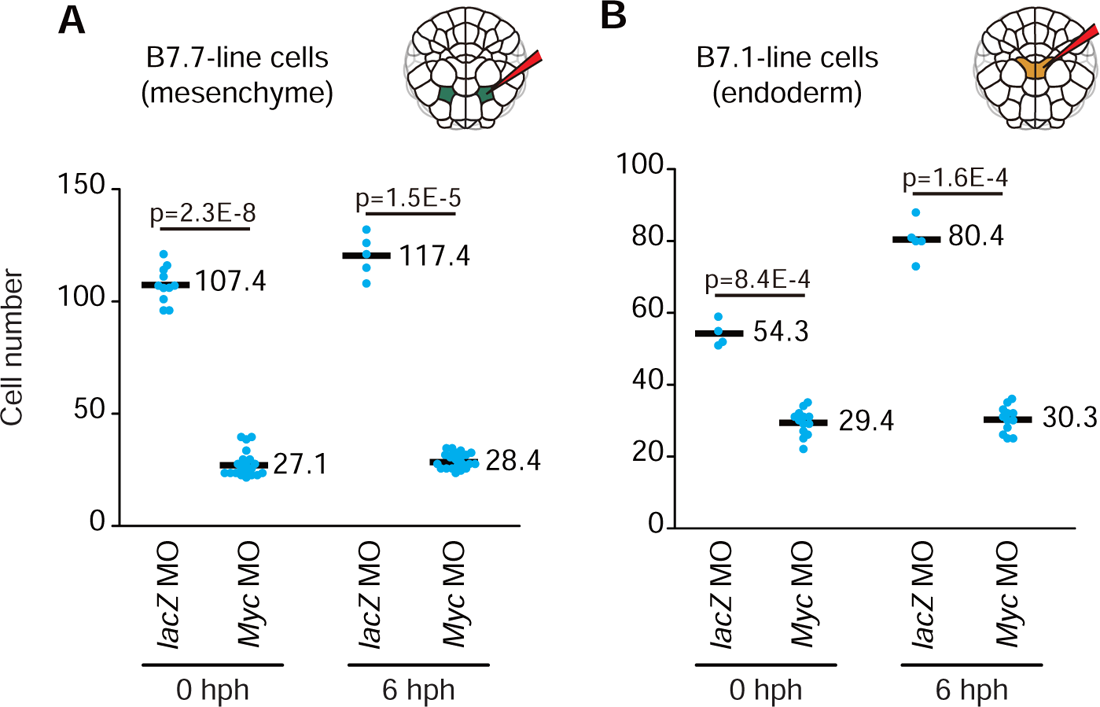
*Myc k*nockdown decreases cell numbers of mesenchymal and endodermal cells. (A, B) The control *lacZ* or *Myc* MO was injected into eggs. At the 64-cell stage, a presumptive mesenchymal cell (B7.7; A) and an endodermal cell (B7.1; B) were labelled with DiI, and incubated until hatching (0 hph) or 6 h after hatching (6 hph). Larvae were fixed and the number of DiI-labelled cells was counted. Each dot indicates the DiI-labelled cell number of a single larva. Differences in cell number between 0 hph and 6 hph were examined with the two-sided Wilcoxon rank sum test, and p-values are indicated. Each dot indicates the number of labelled cells in a single larva, and averages are shown by black bars.

Similarly, when the control MO was injected and B7.1 (presumptive endodermal cell) was labeled with DiI, 54.3 and 80.4 cells were detected at 0 hph and 6 hph (Figure 7B). On the other hand, when the *Myc* MO was injected, 29.4 and 30.3 cells were detected at 0 hph and 6 hph (Figure 7B). We obtained similar results in the other four endodermal lineages (B7.2, A7.1, A7.2 and A7.5; Figure S5B). Therefore, as in the case of mesenchymal cells, our data indicated that *Myc* is necessary for endodermal cells to continue to divide.

We could not obtain another *Myc* MO that could specifically knock-down *Myc*; therefore, we made a construct that drove a dominant negative version of *Myc* (*dnMyc*) under the upstream regulatory region of *Twist-r.a*. Because *C. robusta* (or *C. intestinalis* type A) eggs are more suitable for electroporation, we used *C. robusta* eggs for this experiment. When we introduced a construct that contained a *lacZ* reporter and the same *Twist-r.a* upstream region into fertilized eggs, resultant larvae expressed the reporter in mesenchymal cells. In larvae co-introduced with constructs encoding *dnMyc* and *lacZ*, we found larger cells than those in normal larvae (Figure S6A, B). To quantify larvae with large cells, we first calculated the average area of mesenchymal cells in a photograph of a slice of a normal larva. Next, we picked several cells that looked large in each of 5 control and 8 experimental larvae, and measured areas of confocal images of these cells. As shown in Figure S6C, embryos that expressed *dnMyc* contained larger cells than control embryos. We also counted the number of cells expressing LacZ. The number of LacZ-positive cells was markedly smaller in larvae with *lacZ* and *dnMyc* constructs than in larvae with the *lacZ* construct only (Figure S6D). These results indicated that the dominant negative form of Myc suppressed cell division. These results were consistent with results obtained using the *Myc* MO in *C. savignyi* larvae; therefore, it is likely that the *Myc* MO successfully inhibited translation of *Myc* mRNA.

### *Myc* is under the control of *Twist-r.a/b* and *Lhx3/4* in mesenchymal and endodermal cells

*Twist-r.a/b* and *Lhx3/4* are key transcription factor genes for fate specification of mesenchyme and endoderm, respectively (Imai et al., 2003; Satou et al., 2001b). *Myc* expression was lost in mesenchymal cells of embryos injected with a *Twist-r.a/b* MO and in endodermal cells of embryos injected with an *Lhx3/4* MO (Figure 8). Meanwhile, expression of two mesenchymal marker genes [*Hm13* (previously called *Psl3*) and *Tram1/2*] (Tokuoka et al., 2004) and two endodermal marker genes (*Thr* and *Gata.a*) (Imai et al., 2004; Imai et al., 2006) were not changed in *Myc*-MO-injected embryos (Figure S7). These data indicated that *Myc* is under control of *Twist-r.a/b* and *Lhx3/4* in mesenchymal cells and endodermal cells, respectively, and that *Myc* is not involved in specification of mesenchymal fate or endodermal fate.

**Figure 8.**
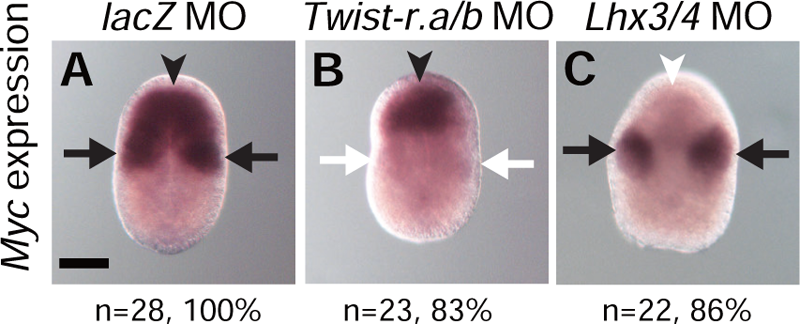
*Myc* is controlled by *Twist-r.a*/b and *Lhx3/4*. *In situ* hybridization of *Myc* at the middle neurula stage in embryos injected with the (A) control *lacZ* MO, (B) *Twist-r.a/b* MO, and (C) *Lhx3/4* MO. Presumptive mesenchymal and endodermal cells are indicated by arrows and arrowheads, respectively. Black arrows and arrowheads indicate expression of *Myc*, and white arrows and arrowheads indicate loss of expression. The number of embryos examined and the proportion of embryos that each photograph represents are shown below the panels. Scale bar, 50 μm.

### Myc regulates transcription of cell cycle regulators

Because Myc is a transcription factor, it may regulate cell cycle regulator genes. To test this hypothesis, we quantified levels of mRNAs encoding Ccna (Cyclin A), Ccnb (Cyclin B), Ccnd (Cyclin D), Ccne (Cyclin E), Cdk1, Cdk2/3, and Cdk4/6 by reverse-transcription followed by quantitative PCR (RT-qPCR). As shown in Figure 9, expression levels of *Ccne*, *Cdk1*, *Cdk2/3*, and *Cdk4/6* were obviously reduced in *Myc-*MO injected embryos, although the difference in *Ccna*, *Ccnb* and *Ccnd* expression levels was not statistically significant.

**Figure 9.**
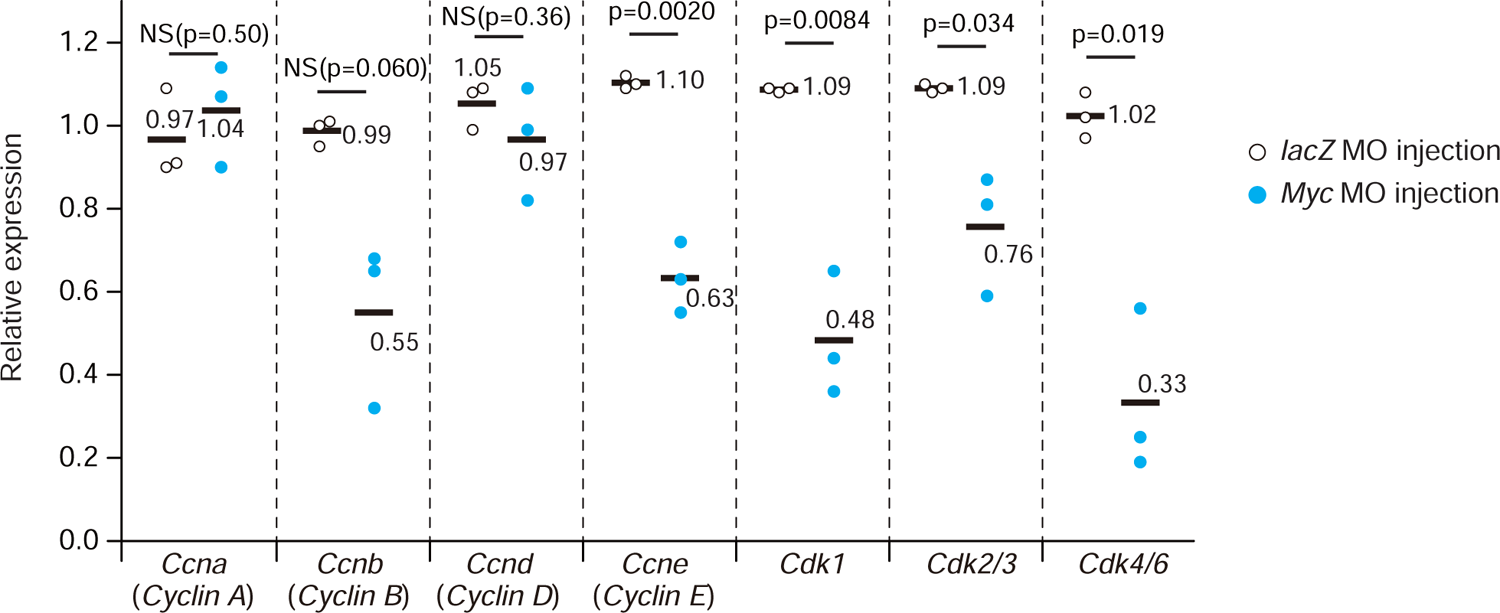
Expression levels of mRNAs encoding cell cycle regulators. Relative mRNA expression levels were measured by RT-qPCR at the larva stage (0 hph). Expression levels in control *lacZ* MO-injected larvae (white dots) and *Myc* MO-injected larvae (cyan dots) are represented as relative values to those of uninjected control larvae. Each dot shows a result for each of three independent experiments. *Ywhaz* was used as an internal control for normalization. Differences in relative expression were analyzed with Student’s t-tests using ΔΔCt values. NS, not statistically significant (p>0.05).

## Discussion

The present study showed that ascidian embryos can support approximately 10 rounds of cell cycles without zygotic transcription. However, numbers of cell cycle rounds in normal embryos are controlled by zygotically expressed *Cdkn1.b* and *Myc*, depending on cell lineages (Figure 10). In the A-line notochord lineage, cells stopped dividing precisely after 9 rounds of cell division under control of *Cdkn1.b*. Similarly, in the primary muscle lineage, cells stopped dividing after 8 (B7.8 lineage) or 9 (B7.4 lineage) rounds of cell division under control of *Cdkn1.b*. Because these numbers are less than the number of cycles that maternal factors can promote, it is reasonable that zygotically expressed *Cdkn1.b* arrests the cell cycle in these tissues.

**Figure 10.**
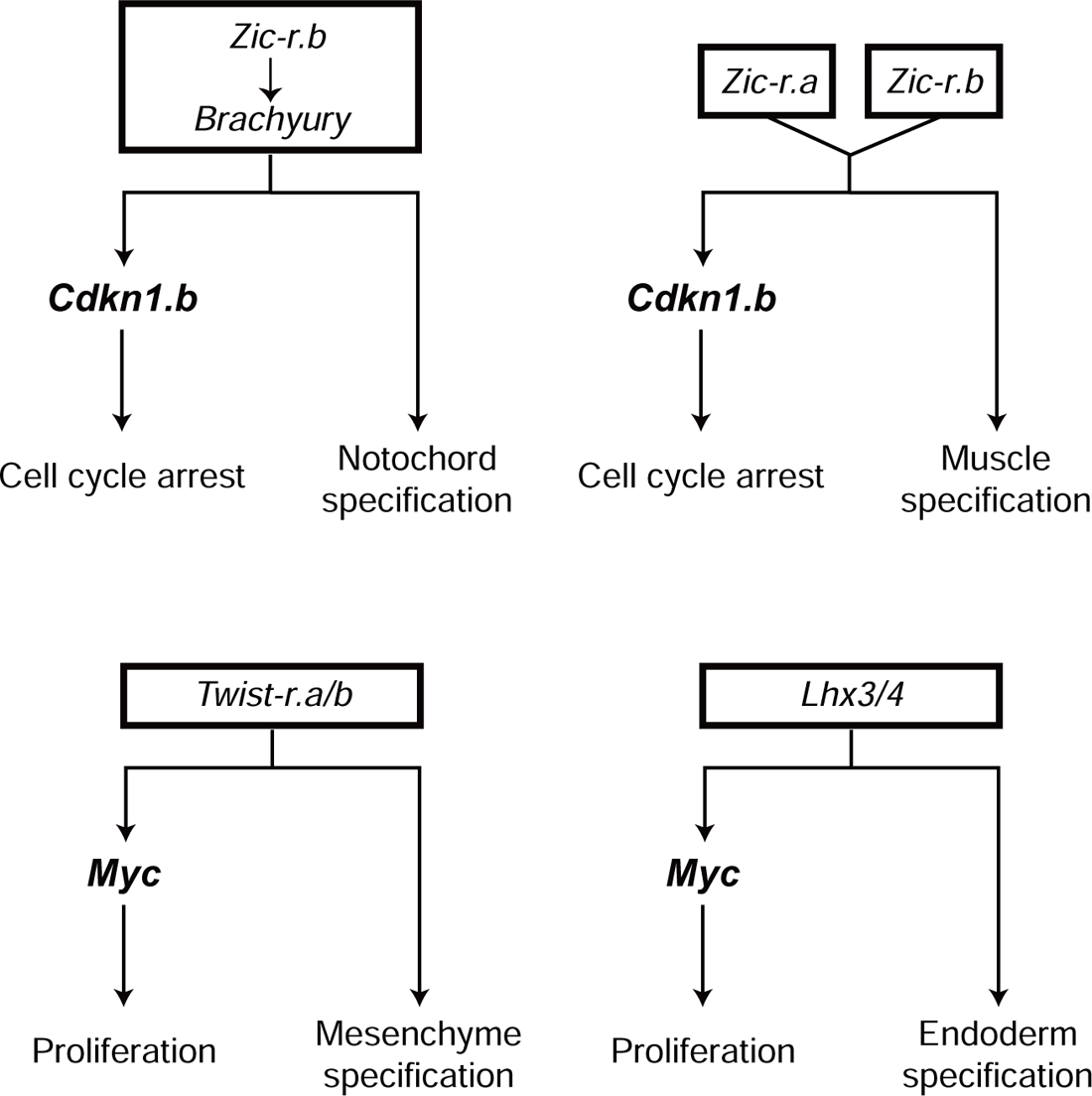
Summary of the present study. In notochord and muscle, cell division is terminated by *Cdkn1.b*, which is activated under control of *Brachyury/Zic-r.b* and *Zic-r.a*/*Zic-r.b,* respectively. In mesenchyme and endoderm, cell division is promoted by *Myc*, which is activated under control of *Twist-r.a/b* and *Lhx3/4,* respectively. *Brachyury*, *Zic-r.a*, *Zic-r.b*, *Twist-r.a/b*, and *Lhx3/4* are key transcription factors for fate specification. In other words, fate specification and cell cycle control are both under control of these transcription factors.

On the other hand, epidermal cells cease cell division after 11 rounds of cell division (Nakayama et al., 2005; Ogura et al., 2011; Pasini et al., 2006). In ascidian embryos, maternal Gata.a specifically activates the developmental program for differentiation of epidermis, and Gata.a activity is restricted to the animal hemisphere, which gives rise to ectodermal tissues, in early embryos (Bertrand et al., 2003; Oda-Ishii et al., 2016; Rothbächer et al., 2007). It is possible that epidermal progenitors also contain more abundant maternal products promoting cell cycle than other cells in early embryos. Even if so, as in the cases of notochord and muscle, *Cdkn1.b* ensured that epidermal cells stop dividing after 11 rounds of cell division. It is also possible that a factor promoting cell division other than Myc is expressed before *Cdkn1.b* expression.

Mesenchymal and endodermal cells do not have apparent functions in larvae, and they produce adult mesodermal and endodermal tissues after metamorphosis. The number of mesenchymal cells is estimated to be around 900 at 0 hph (Monroy, 1979). As we showed in the present study, numbers of the three lines of mesenchymal cells became 1.1–1.6 times (1.3 times on average) larger at 6 hph than at 0 hph (Figure S3); therefore, there may be ∼1200 cells at 6 hph. Because mesenchymal cells are derived from five pairs of cells (10 cells) in the eighth generation (Nishida, 1987), these presumptive cells are estimated to divide 6 to 7 more times (13 to 14 times in total from 1-cell embryos).

Similarly, the number of endodermal cells is estimated to be around 500 at 0 hph (Monroy, 1979). We observed that numbers of the five lineages of endodermal cells became 1.1–1.9 times (1.6 times on average) greater at 6 hph than at 0 hph; therefore, there may be ∼800 cells at 6 hph. Endodermal cells are derived from five pairs of cells in the seventh generation. Hence, it is likely that the five pairs of presumptive endodermal cells (10 cells) divide 6 to 7 additional times (12 to 13 times in total).

Thus, numbers of cell cycles of mesenchymal and endodermal cells greatly exceed the number of cell cycles that maternal factors can promote. The present study indicated that *Myc* expression is required for mesenchymal and endodermal cells to divide so many times probably through activating *Ccne*, *Cdk1*, *Cdk2/3*, and *Cdk4/6*.

*Myc* is a well-known oncogene that controls cell cycling. One function of *Myc* is to regulate cell cycle-related genes (Bretones et al., 2015). Therefore, the ascidian uses this conserved mechanism to continue the cell cycle in embryonic/larval mesenchyme and endoderm. On the other hand, Myc also regulates the cell cycle by hyperactivating Cyclin/Cdk complexes, inhibiting transcription of *Cdkn* encoding a Cdk inhibitor, and activating genes that encode proteins involved in DNA replication (Bretones et al., 2015). It is possible that these mechanisms also act in ascidian mesenchyme and endoderm.

Inhibition of progression of the cell cycle by Cdk inhibitors and maintenance of cell proliferation by Myc are well known mechanisms for cell cycle control in animal development (deNooij et al., 1996; Fukuyama et al., 2003; Knoepfler et al., 2002; Lane et al., 1996; Liu et al., 2012). The present study showed that both of these mechanisms are used in different tissues of ascidian larvae, and identified their upstream regulators; *Cdkn1.b* is under control of *Brachyury* (and its upstream regulator *Zic-r.b*) in the notochord lineage and under control of *Zic-r.a* and *Zic-r.b* in the muscle lineage, while *Myc* is controlled by *Twist-r.a/b* in the mesenchymal lineage and by *Lhx3/4* in the endodermal lineage. These upstream regulators are transcription factors important for specifying cell fate (Corbo et al., 1997; Imai et al., 2003; Imai et al., 2002; Nishida and Sawada, 2001; Reeves et al., 2017; Satou et al., 2001b; Takahashi et al., 1999; Yasuo and Satoh, 1993; Yasuo and Satoh, 1994; Yu et al., 2019). In muscle cells, *Tbx6-r.b* is activated under control of *Zic-r.a* and *Zic-r.b* (Yagi et al., 2004; Yagi et al., 2005; Yu et al., 2019), and involvement of *Tbx6-r.b* in cell cycle regulation in muscle cells has been suggested in *Halocynthia* embryos (Kuwajima et al., 2014). Therefore, it is possible that *Cdkn1.b* is under control of *Tbx6-r.b* in *Ciona* embryos. Therefore, the present study showed that the same transcription factors that specify cell fate also regulate cell cycle regulator genes. In a related species, *Ciona robusta* (*Ciona intestinalis* type A), *Dlx.b* and *Tfap2-r.b* contribute to epidermal fate specification (Imai et al., 2017). Although currently, we cannot efficiently knock-down or knock-out these genes in *C. savignyi*, it is possible that *Cdkn1.b* is regulated by either or both of these factors. Thus, cell fate specification and cell cycle control are tightly linked at the point of these key transcription factors in ascidian embryos.

## Materials and Methods

### Animals and gene identifiers

Adult specimens of *Ciona savignyi* were obtained from International Coastal Research Center (the University of Tokyo), Onagawa Field Center (Tohoku University), Research Center for Marine Biology (Tohoku University), Maizuru Fisheries Research Station (Kyoto University), Honmoku fishing harbor (Kanagawa, Japan), and University of California, Santa Barbara (USA). Identifiers in Ensembl (Howe et al., 2021) for genes examined in this study are ENSCSAVG00000005700 for *Cdkn1.b*, ENSCSAVG00000000363 for *Myc*, ENSCSAVG00000002247.1 for *Brachyury*, ENSCSAVG00000001907 for *Zic-r.a*, ENSCSAVG00000002442/2450/2454/2459/2465 for *Zic-r.b* (multiple copies exist), ENSCSAVG00000011698 for *Talin*, ENSCSAVG00000009518 for *Tropomyosin-like 1*, ENSCSAVG00000008180 for *Ma1*, ENSCSAVG00000003085 for *Epi1*, ENSCSAVG00000009525 for *Cesa*, ENSCSAVG00000008611/8613for *Twist-r.a/b*, ENSCSAVG00000000590 for *Lhx3/4*, ENSCSAVG00000007855 for *Hm13* (*Psl3*), ENSCSAVG00000009046 for *Tram1/2*, ENSCSAVG00000006056 for *Thr*, ENSCSAVG00000008350 for *Gata.a*, ENSCSAVG00000010339 for *Ccna* (*Cyclin A)*, ENSCSAVG00000003185 for *Ccnb* (*Cyclin B*), ENSCSAVG00000003746 for *Ccnd* (*Cyclin D*), ENSCSAVG00000005373 for *Ccne* (*Cyclin E*), ENSCSAVG00000001660 for *Cdk1*, ENSCSAVG00000009914 for *Cdk2/3*, and ENSCSAVG00000005088 for *Cdk4/6*.

Adult specimens of *Ciona robusta* (also called *Ciona intestinalis* type A) were obtained from the National Bio-Resource Project for *Ciona*. Identifiers for genes examined in this study are KY21.Chr1.384 for *Myc* and KY21.Chr5.361 for *Twist-r.a* [gene models were based on the annotation for the latest genome assembly of this animal (Satou et al., 2019; Satou et al., 2022)].

### Gene knockdown and over/ectopic expression of *Myc*

Nucleotide sequences of MOs (custom made by Gene Tools) were as follows; *Cdkn1.b*, 5’-CATTGTACGAAGGCGGTGGAACCAT-3’, and 5’-AAGGCGGTGGAACCATTTTGATTAC-3’; *Myc*, 5ʹ-TGTTTATCGGTGTGCTGAGCTTCAT-3ʹ; *Brachyury*, 5ʹ-AACACGATTCTAAAGTGCTGGTCAT-3ʹ; *Twist-r.a/b*, 5ʹ-CTTGATTGTACTCTAGTGATGTCAT-3ʹ; *Lhx3/4*, 5ʹ-AAAGCGGGCTTGACTCGATACACAT-3ʹ; *Zic-r.a*, 5ʹ-AACCAAGCGTGCCAACGAAAGCCAT-3ʹ; *Zic-r.b*, 5ʹ-GTACATGATATTGATTGCTGTCTAA-3ʹ. These MOs were designed to block translation. MOs for *Brachyury*, *Twist-r.a/b*, *Zic-r.a*, and *Zic-r.b* were previously used. Specificity of MOs for *Cdkn1.b* and *Myc* was examined and is described in the Results section. In larvae injected with the *Lhx3/4* MO, alkaline phosphatase activity was rarely detected by histochemical staining (Figure S8). This phenotype was similar to that obtained in a previous study in which a different MO against *Lhx3/4* was used, suggesting that the *Lhx3/4* MO we used in the present study acted specifically (Satou et al., 2001b). We also used an MO against *Escherichia coli lacZ* as a negative control (5ʹ-TACGCTTCTTCTTTGGAGCAGTCAT-3ʹ). These MOs were microinjected into unfertilized eggs unless otherwise specified. The concentration of the MOs was 1.5 mM for *Cdkn1.b*, 1.0 mM for *Brachyury* and *Zic-r.b*, and 0.5 mM for the others.

A mutant *C. robusta Myc* that encoded a dominant negative form of Myc lacking Myc box II (Haque et al., 2016) was fused to the upstream region of *Twist-r.a*, and *E. coli lacZ* was similarly fused to the upstream region of *Twist-r.a*. These constructs were introduced into *C. robusta* fertilized eggs by electroporation. To measure the area of a cell, we used a confocal slice in which a given cell looked largest among all slices, and approximated each cell using polygons.

### Isolation of blastomeres, DiI labelling, cell count, and inhibition of transcription

Blastomeres were isolated with a fine glass needle under a binocular microscope, and isolated blastomeres were incubated using agar-coated dishes. DiI (CellTracker CM-DiI, Thermo Fisher Scientific) was dissolved in soybean oil at a concentration of 1 mg/mL. We labeled blastomeres with the DiI solution under a binocular microscope. To count cell numbers, embryos and larvae were fixed for 30 min in 4% paraformaldehyde solution and stained with DAPI (4’,6-diamidino-2-phenylindole). Numbers of nuclei were counted under a fluorescence microscope or a confocal laser-scanning microscope. α-amanitin was injected into unfertilized eggs at the concentration of 2 μg/mL. Actinomycin D was dissolved in DMSO and added to sea water at a final concentration of 200 μg/mL.

### Whole mount *in situ* hybridization, phalloidin staining, and histochemical staining for alkaline phosphatase and acetylcholine esterase

Whole-mount *in situ* hybridization was performed as described previously (Satou et al., 1995). To delineate cell shape, embryos were stained with Alexa Fluor 488 Phalloidin (Thermo Fisher Scientific, A12379) after fixation with 4% paraformaldehyde in MOPS buffer (0.5 mM NaCl, 0.1 M MOPS, pH7.5).

For histochemical detection of alkaline phosphatase activity, embryos were fixed with 4% paraformaldehyde in MOPS buffer (0.5 mM NaCl, 0.1 M MOPS, pH7.5) for 10 min at room temperature. Embryos were washed in AP staining buffer (100 mM NaCl, 50 mM MgCl_2_, 100 mM Tris-HCl, pH 9.5) twice, and then incubated with staining buffer (2.25 mg/mL Nitro Blue Tetrazolium, 1.75 mg/ml 5-bromo-4-chloro-3-indolyl phosphate).

For histochemical detection of acetylcholinesterase (AChE), embryos were fixed with 4% paraformaldehyde in MOPS buffer (0.5 mM NaCl, 0.1 M MOPS, pH 7.5) for 10 min at room temperature. Fixed embryos were washed in 100 mM sodium phosphate buffer three times, and incubated in AChE staining buffer (0.5 mg/mL acetylthiocholine, 65 mM sodium phosphate, 3 mM copper sulfate, 0.5 mM potassium ferricyanide, 5 mM sodium citrate, pH 6.0).

### BrdU incorporation and immunodetection

To monitor DNA synthesis, we added BrdU (Nacalai Tesque, 05650) to sea water at a concentration of 100 μM. Incorporation of BrdU was monitored with a specific antibody (Merck, 11170376001) following a protocol described previously (Nakayama et al., 2005).

### Reverse transcription and quantitative PCR

Total RNA was extracted from 30 larvae (0 hph) using NucleoSpin RNA XS (Macherey-Nagel, U0902A). cDNA was synthesized with an iScript cDNA synthesis kit (Bio-Rad, 1708891). Quantitative PCR was performed using the MiniOpticon Real-Time PCR Detection System (Bio-Rad) with SsoFast Evagreen Supermix (Bio-Rad, 1725200). A one-embryo-equivalent quantity of cDNA was used for each reaction. Cycling conditions were preheating at 95 °C for 30 s, and 40 cycles of 95 °C for 5 s and 60 °C for 10 s. We took three biological replicates. Relative expression values were calculated using the ΔΔCT method, and expression values were normalized to a housekeeping gene, *Ywhaz* (tyrosine 3-monooxygenase/tryptophan 5-monooxygenase activation protein, zeta polypeptide). *Ccna*, 5ʹ-GCAGCAAGACATCACAGTTGG-3ʹ and 5ʹ-TGGTTTCGGTGTGGAGTTTG-3ʹ; *Ccnb*, 5ʹ-CAACATACGCTCGCAAAATACC-3ʹ and 5ʹ-AAGGCACAAGGAACCAGCA-3ʹ; *Ccnd*, 5ʹ-AAGGTGTGAAGATGATGTGTTTCC-3ʹ and 5ʹ-GTTGAGTTCGTTGGATTGGTTG-3ʹ; *Ccne*, 5ʹ-GGAGGTGTGCGAGGTTTATTC-3ʹ and 5ʹ-GGGTTTTATGGATGTCGGTTG-3ʹ; *Cdk1*, 5ʹ-CGGTGTGACGCAGTTGAAAG-3ʹ and 5ʹ-ATACGAGGCATTTAGCGAGCA-3ʹ; *Cdk2/3,* 5ʹ-TCGCACAGAGTCCTACACAGAGA-3ʹ and 5ʹ-ATACATCCGCACTGGCACAC-3ʹ; *Cdk4/6*, 5ʹ-AGCGTGAAACCCAACTAATGC-3ʹ and 5ʹ-AGTCCCATCAACATCTGCCTC-3ʹ. *Ywhaz*, 5’-TTTCTGGACTTGGACGCAAAC-3’ and 5’-CGGCTTTCCCTTGTCCTTATC-3’

## Acknowledgements

We thank Masatoshi Hara and Sawako Hori-Oshima for earlier works. We are greatful to the staff of International Coastal Research Center (the University of Tokyo), Onagawa Field Center (Tohoku University), Research Center for Marine Biology (Tohoku University), Maizuru Fisheries Research Station (Kyoto University) and Honmoku fishing harbor (Kanagawa, Japan) for their help in collecting *Ciona savignyi* adults. We also thank Shota Chiba, Erin Newman-Smith, and William C. Smith at University of California, Santa Barbara (USA) for providing *Ciona savignyi* adults. We thank Manabu Yoshida (the University of Tokyo) and other members working under the National Bio-Resource Project for *Ciona* (MEXT, Japan) at Kyoto University and the University of Tokyo for providing *Ciona robusta* adults. A draft of the manuscript was edited by a technical editor, Steven D. Arid.

---

**Figure S1.**
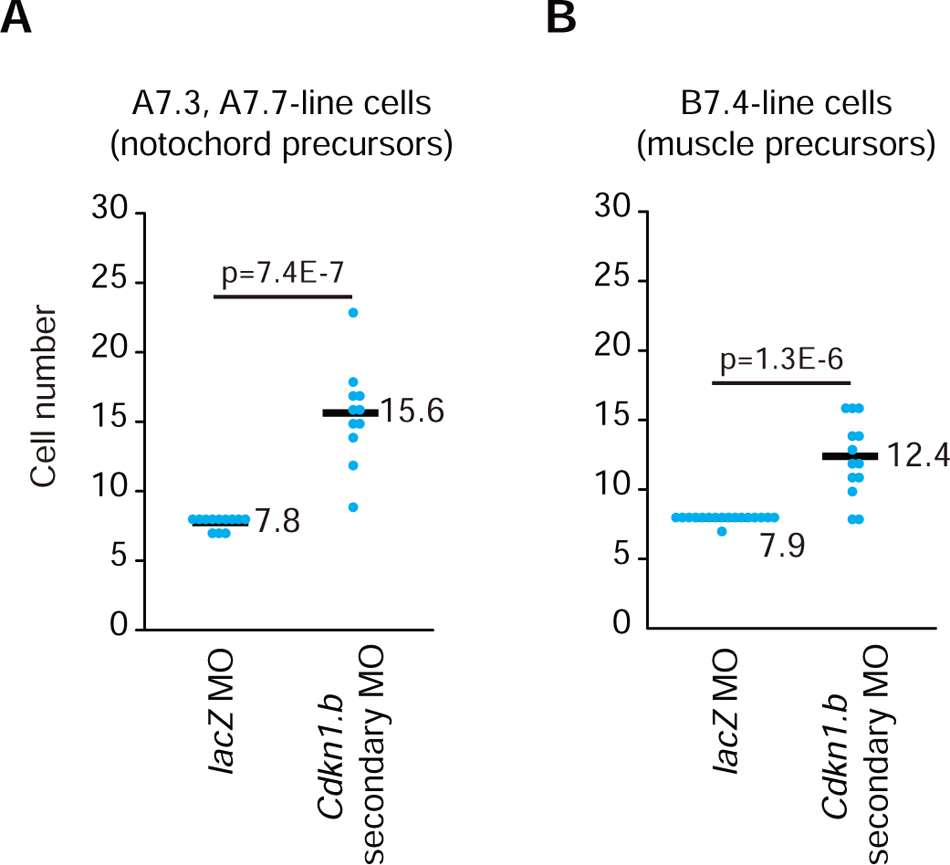
A second MO against *Cdkn1.b* gave the same phenotype as the first MO. Presumptive notochord, or muscle cells isolated from embryos injected with the second *Cdkn1.b* MO produced larger numbers of cells than those isolated from control embryos. The control *lacZ* or *Cdkn1.b* MO was injected into eggs. At the 64-cell stage, a presumptive notochord (A) or muscle (B; B7.4) cell was isolated and incubated until uninjected sibling embryos hatched (0 hph). Embryos were fixed and cell numbers were counted. Differences in cell number were examined with the two-sided Wilcoxon rank sum test, and p-values are indicated. Each dot indicates the number of labelled cells in a single embryo, and averages are shown by black bars.

**Figure S2.**
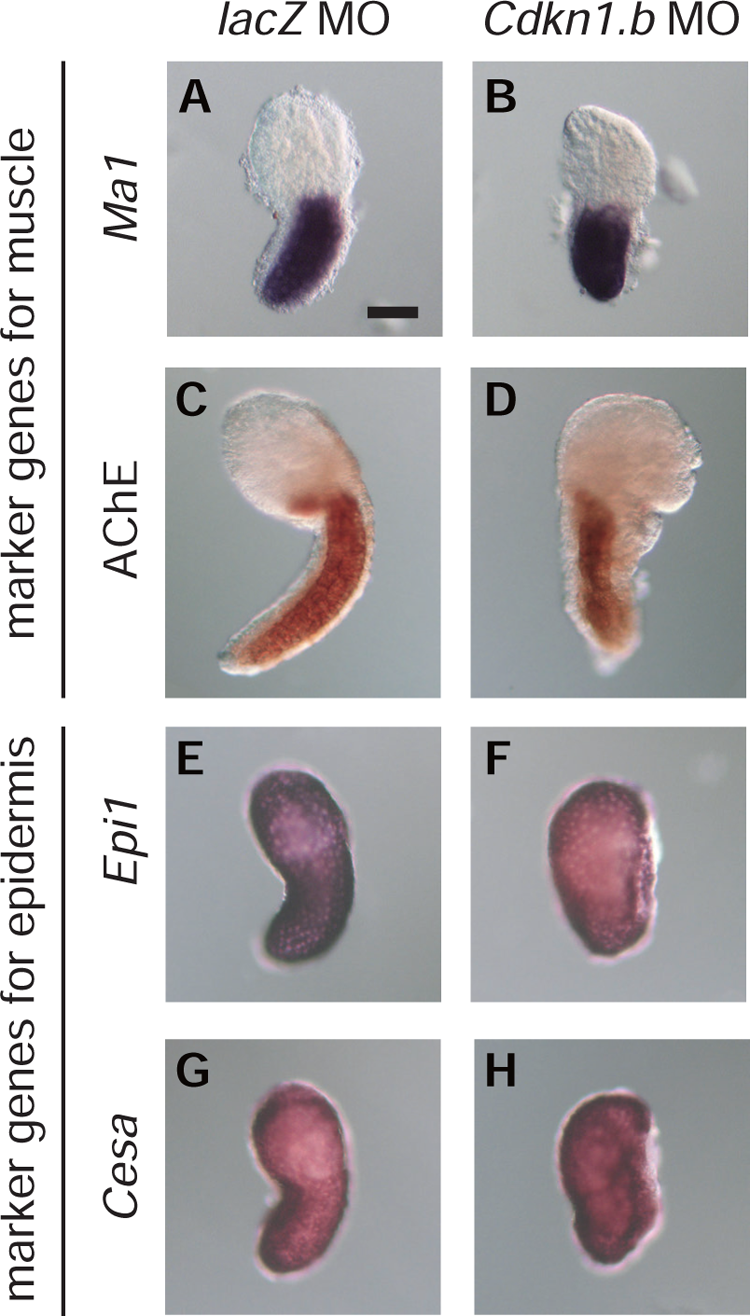
Differentiation markers of muscle and epidermis. Differentiation markers for muscle (A-D) and epidermis (E-H) were tested in embryos injected with the control *lacZ* MO (A, C, E, G) or *Cdkn1.b* MO (B, D, F, H). Expression of (A, B) muscle actin gene (*Ma1*), (E, F) *Epi1*, and (G, H) *Cesa* was examined by *in situ* hybridization. (C, D) Expression of acetylcholine esterase (ACHE) was examined by histochemical staining. Scale bar, 50 μm.

**Figure S3.**
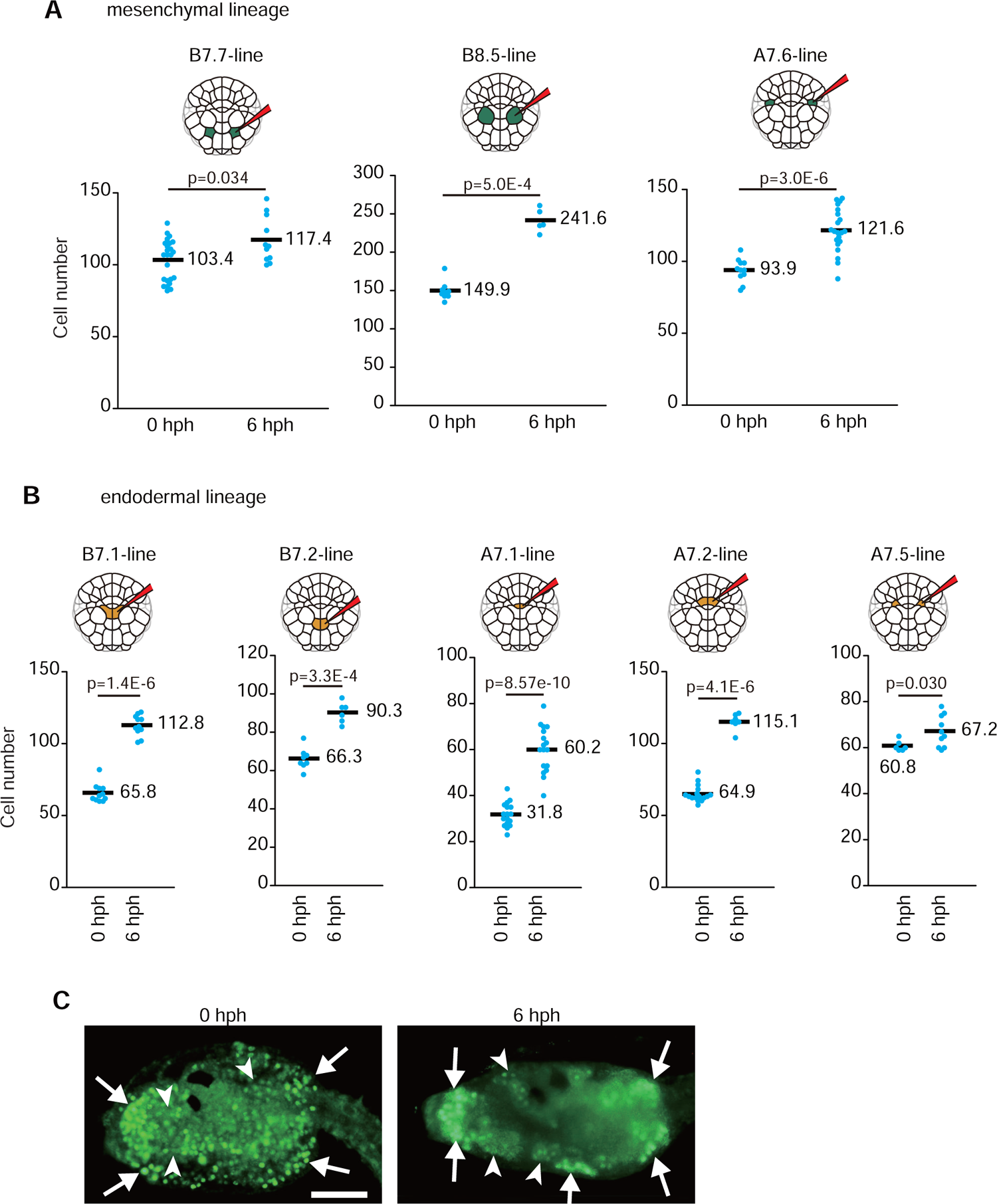
Mesenchymal and endodermal cells divide even after hatching. (A, B) At the 112-cell stage, one of three presumptive mesenchymal cells (A) or one of five presumptive endodermal cells (B) was labelled with DiI. Numbers of labelled cells were counted immediately after hatching (0 hph) or 6 h later (6 hph). Differences in cell number between 0 hph and 6 hph were examined with the one-sided Wilcoxon rank sum test, and p-values are indicated. Each dot indicates the number of labelled cells in a single embryo, and averages are shown by black bars. (C) Larval trunk regions stained with an anti-BrdU antibody. This larva was incubated in sea water containing BrdU for 30 min from 0 hph (left) and 6 hph (right). Signals are evident in mesenchymal and endodermal cells. Some mesenchymal cells and endodermal cells are indicted by arrows and arrowheads, respectively. Scale bar, 50 μm.

**Figure S4.**
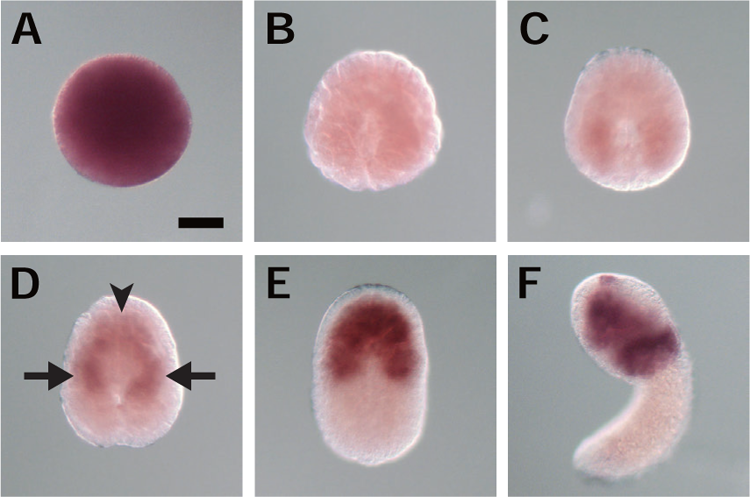
Expression pattern of *Myc* in *C. savignyi* embryos. Photographs of whole mount *in situ* hybridization of (A) a fertilized egg, and embryos at the (B) middle gastrula, (C) late gastrula, (D) early neurula, (E) middle neurula, and (F) early tailbud stages. Expression in the mesenchyme lineage was first evident at the early neurula stage (arrows). Expression in the endoderm lineage was also first evident at the early neurula stage (arrowhead). Scale bar, 50 μm.

**Figure S5.**
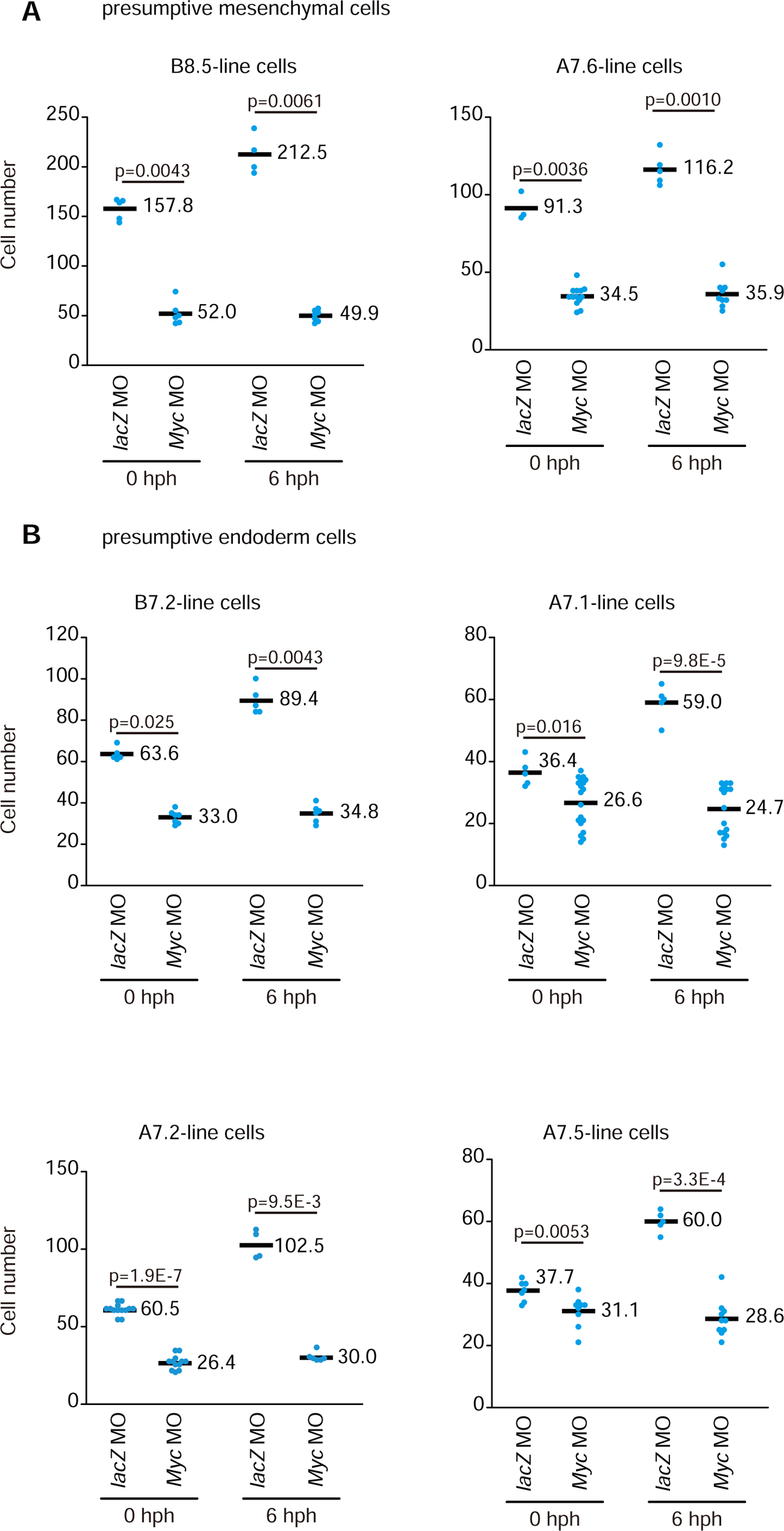
Knockdown of *Myc* decreases numbers of mesenchymal and endodermal cells. **(A,** B) The control *lacZ* or *Myc* MO was injected into eggs. At the 64-cell stage, a presumptive mesenchymal cell (B8.5 or A7.6; A) and endodermal cell (B7.2, A7.1, A7.2, or A7.5; B) was labelled with DiI, and incubated until hatching (0 hph) or 6 h after hatching (6 hph). Larvae were fixed and numbers of DiI-labelled cells were counted. Differences in cell number between larvae injected with the control *lacZ* MO and *Myc* MO were examined with the two-sided Wilcoxon rank sum test, and p-values are indicated. Each dot indicates the number of labelled cells in a single larva, and averages are shown by black bars.

**Figure S6.**
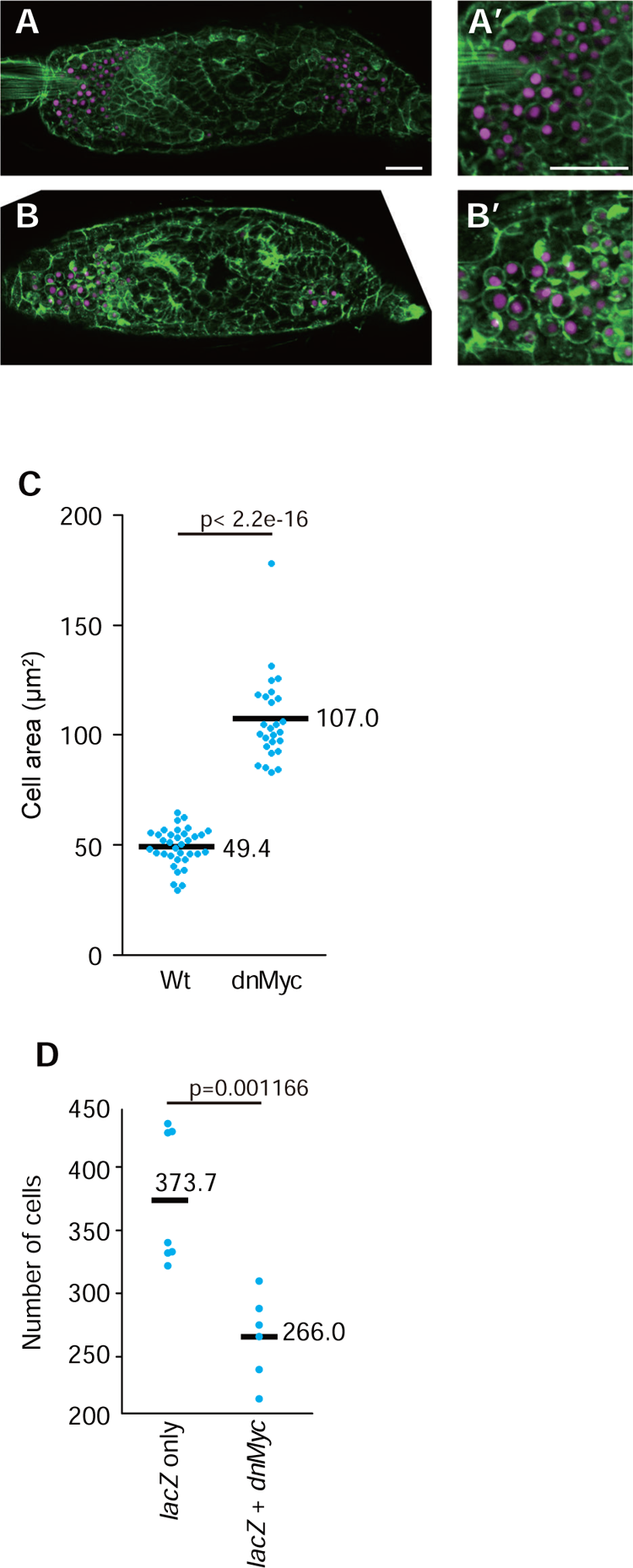
A dominant negative form of *Myc* suppresses cell division in the mesenchymal and endodermal lineages. (A, B) A DNA construct that expressed *lacZ* under control of the upstream regulatory sequence of *Twist-r.a* was introduced into fertilized eggs of *C. robusta* (*C. intestinalis* type A) by electroporation. In B, a DNA construct that expressed a dominant negative form of Myc under the same upstream sequence was co-introduced. Larvae were fixed at 6 hph. An anti-lacZ antibody was used to detect lacZ protein (magenta). Larvae were also stained with phalloidin (green). Higher magnification views of trunk regions close to the tail are shown (A’, B’). Scale bars, 25 μm. (C) We measured cell areas in confocal slices of 35 cells from five control embryos, and 26 cells from eight embryos introduced with *dnMyc*. Embryos were fixed at the late tailbud stage. We picked several cells that looked large in individual embryos. Each dot represents a cell, and mean values are shown by black bars. Differences in cell area were examined with the two-sided Wilcoxon rank sum test, and p-values are indicated. (D) Numbers of lacZ-positive cells were smaller in larvae that expressed lacZ and a dominant negative from of Myc than in larvae that expressed lacZ only. Larvae were fixed at 6 hph. The difference in cell number was examined with the two-sided Wilcoxon rank sum test, and p-values are indicated. Each dot indicates the number of labelled cells in a single larva, and mean values are shown by black bars.

**Figure S7.**
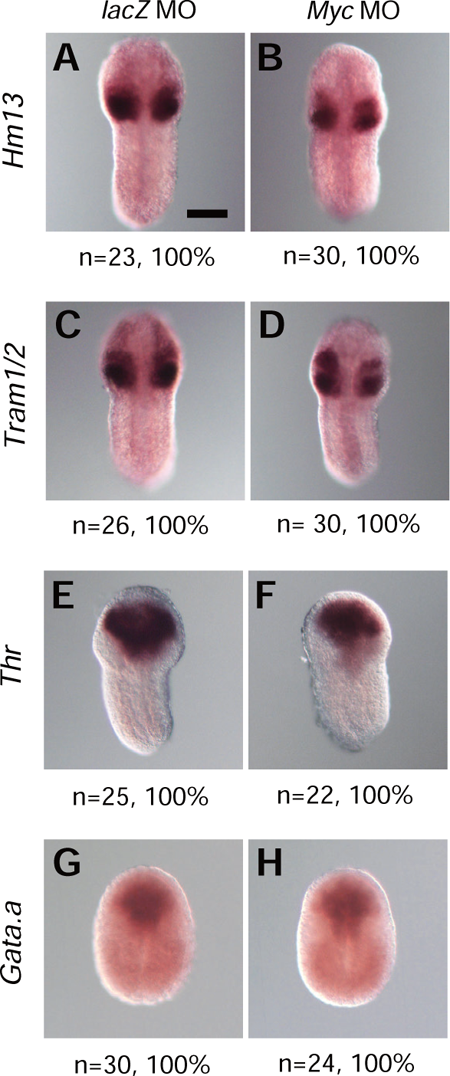
Expression of marker genes for mesenchymal and endodermal cells. Expression of two mesenchymal marker genes, (A, B) *Hm13* and (C, D) *Tram1/2*, and two endodermal marker genes, (E, F) *Thr* and (G, H) *Gata.a*, was examined by *in situ* hybridization at (A-F) the early tailbud stage and (G, H) the middle neurula stage in (A, C, E, G) control *lacZ* MO-injected embryos and (B, D, F, H) *Myc* MO-injected embryos. Numbers of embryos examined and proportions of embryos that each photograph represents are shown below the panels. Scale bar, 50 μm.

**Figure S8.**
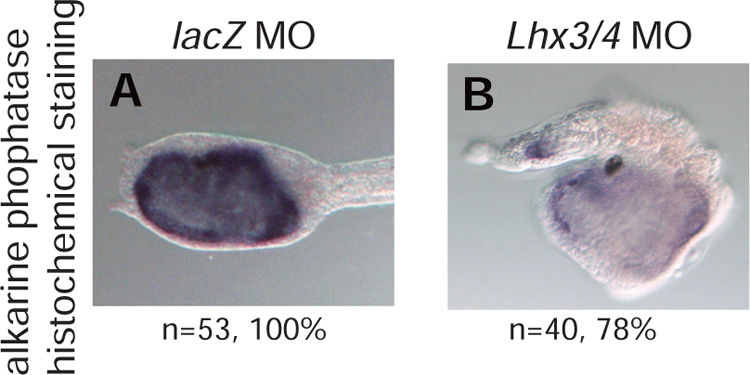
Detection of alkaline phosphatase activity by histochemical staining. Larvae developed from eggs injected with (A) the control *lacZ* MO and (B) the *Lhx3/4* MO were histochemically stained to detect alkaline phosphatase activity. Numbers of larvae examined and proportions of larvae that each photograph represents are shown below the panels. In A, only the trunk region was shown. Loss of alkaline phosphatase activity has been reported in a previous study that used a different MO against *Lhx3/4* (Satou et al., 2001b). Scale bar, 50 μm.

## References

1. Bertrand, V., Hudson, C., Caillol, D., Popovici, C. and Lemaire, P. (2003). Neural tissue in ascidian embryos is induced by FGF9/16/20, acting via a combination of maternal GATA and Ets transcription factors. Cell 115, 615–627.

2. Bretones, G., Delgado, M. D. and Leon, J. (2015). Myc and cell cycle control. Biochimica Et Biophysica Acta-Gene Regulatory Mechanisms 1849, 506–516.

3. Chiba, S., Jiang, D., Satoh, N. and Smith, W. C. (2009). Brachyury null mutant-induced defects in juvenile ascidian endodermal organs. Development 136, 35–39.

4. Conklin, E. G. (1905). The Organization and Cell-Lineage of the Ascidian Egg. *J. Acad.*, Nat. Sci. Phila. 13, 1–119.

5. Corbo, J. C., Levine, M. and Zeller, R. W. (1997). Characterization of a notochord-specific enhancer from the Brachyury promoter region of the ascidian, Ciona intestinalis. Development 124, 589–602.

6. deNooij, J. C., Letendre, M. A. and Hariharan, I. K. (1996). A cyclin-dependent kinase inhibitor, dacapo, is necessary for timely exit from the cell cycle during Drosophila embryogenesis. Cell 87, 1237–1247.

7. Di Gregorio, A. and Levine, M. (1999). Regulation of Ci-tropomyosin-like, a Brachyury target gene in the ascidian, Ciona intestinalis. Development 126, 5599–5609.

8. Fujikawa, T., Takatori, N., Kuwajima, M., Kim, G. J. and Nishida, H. (2011). Tissue-specific regulation of the number of cell division rounds by inductive cell interaction and transcription factors during ascidian embryogenesis. Developmental Biology 355, 313–323.

9. Fukuyama, M., Gendreau, S. B., Derry, W. B. and Rothman, J. H. (2003). Essential embryonic roles of the CKI-1 cyclin-dependent kinase inhibitor in cell-cycle exit and morphogenesis in C-elegans. Developmental Biology 260, 273–286.

10. Haque, M., Song, J., Fino, K., Wang, Y., Sandhu, P., Song, X., Norbury, C., Ni, B., Fang, D., Salek-Ardakani, S., et al. (2016). C-Myc regulation by costimulatory signals modulates the generation of CD8+ memory T cells during viral infection. Open Biol 6, 150208.

11. Howe, K. L., Achuthan, P., Allen, J., Allen, J., Alvarez-Jarreta, J., Amode, M. R., Armean, I. M., Azov, A. G., Bennett, R., Bhai, J., et al. (2021). Ensembl 2021. Nucleic Acids Research 49, D884–D891.

12. Imai, K., Satoh, N. and Satou, Y. (2003). A Twist-like bHLH gene is a downstream factor of an endogenous FGF and determines mesenchymal fate in the ascidian embryos. Development 130, 4461–4472.

13. Imai, K. S., Hikawa, H., Kobayashi, K. and Satou, Y. (2017). *Tfap2* and *Sox1/2/3* cooperatively specify ectodermal fates in ascidian embryos. Development 144, 33–37.

14. Imai, K. S., Hino, K., Yagi, K., Satoh, N. and Satou, Y. (2004). Gene expression profiles of transcription factors and signaling molecules in the ascidian embryo: towards a comprehensive understanding of gene networks. Development 131, 4047–4058.

15. Imai, K. S., Levine, M., Satoh, N. and Satou, Y. (2006). Regulatory blueprint for a chordate embryo. Science 312, 1183–1187.

16. Imai, K. S., Satou, Y. and Satoh, N. (2002). Multiple functions of a Zic-like gene in the differentiation of notochord, central nervous system and muscle in Ciona savignyi embryos. Development 129, 2723–2732.

17. Jiang, D. and Smith, W. C. (2007). Ascidian notochord morphogenesis. Dev Dyn 236, 1748–1757.

18. Knoepfler, P. S., Cheng, P. F. and Eisenman, R. N. (2002). N-myc is essential during neurogenesis for the rapid expansion of progenitor cell populations and the inhibition of neuronal differentiation. Genes & Development 16, 2699–2712.

19. Kuwajima, M., Kumano, G. and Nishida, H. (2014). Regulation of the Number of Cell Division Rounds by Tissue-Specific Transcription Factors and Cdk Inhibitor during Ascidian Embryogenesis. Plos One 9.

20. Lane, M. E., Sauer, K., Wallace, K., Jan, Y. N., Lehner, C. F. and Vaessin, H. (1996). Dacapo, a cyclin-dependent kinase inhibitor, stops cell proliferation during Drosophila development. Cell 87, 1225–1235.

21. Liu, Q. C., Zha, X. H., Faralli, H., Yin, H., Louis-Jeune, C., Perdiguero, E., Pranckeviciene, E., Munoz-Canoves, P., Rudnicki, M. A., Brand, M., et al. (2012). Comparative expression profiling identifies differential roles for Myogenin and p38 alpha MAPK signaling in myogenesis. J Mol Cell Biol 4, 386–397.

22. Monroy, A. (1979). Introductory remarks on the segregation of cell lines in the embryo. In Cell lineage, stem cells and cell determination (ed. N. L. Douarin), pp. 3-13: Elsevier/North-Holland Biomedical Press.

23. Nakayama, A., Satoh, N. and Sasakura, Y. (2005). Tissue-specific profile of DNA replication in the swimming larvae of Ciona intestinalis. Zool Sci 22, 301–309.

24. Nishida, H. (1987). Cell lineage analysis in ascidian embryos by intracellular injection of a tracer enzyme. III. Up to the tissue restricted stage. Dev Biol 121, 526–541.

25. Nishida, H. and Sawada, K. (2001). macho-1 encodes a localized mRNA in ascidian eggs that specifies muscle fate during embryogenesis. Nature 409, 724–729.

26. Oda-Ishii, I., Kubo, A., Kari, W., Suzuki, N., Rothbacher, U. and Satou, Y. (2016). A maternal system initiating the zygotic developmental program through combinatorial repression in the ascidian embryo. PLoS genetics 12, e1006045.

27. Ogura, Y., Sakaue-Sawano, A., Nakagawa, M., Satoh, N., Miyawaki, A. and Sasakura, Y. (2011). Coordination of mitosis and morphogenesis: role of a prolonged G2 phase during chordate neurulation. Development 138, 577–587.

28. Pasini, A., Amiel, A., Rothbacher, U., Roure, A., Lemaire, P. and Darras, S. (2006). Formation of the ascidian epidermal sensory neurons: insights into the origin of the chordate peripheral nervous system. PLoS Biol 4, e225.

29. Reeves, W. M., Wu, Y. Y., Harder, M. J. and Veeman, M. T. (2017). Functional and evolutionary insights from the Ciona notochord transcriptome. Development 144, 3375–3387.

30. Rothbächer, U., Bertrand, V., Lamy, C. and Lemaire, P. (2007). A combinatorial code of maternal GATA, Ets and β-catenin-TCF transcription factors specifies and patterns the early ascidian ectoderm. Development 134, 4023–4032.

31. Satou, Y., Imai, K. and Satoh, N. (2001a). Action of morpholinos in Ciona embryos. Genesis 30, 103–106.

32. Satou, Y., Imai, K. S. and Satoh, N. (2001b). Early embryonic expression of a LIM-homeobox gene *Cs-lhx3* is downstream of β-catenin and responsible for the endoderm differentiation in *Ciona savignyi* embryos. Development 128, 3559–3570.

33. Satou, Y., Kusakabe, T., Araki, S. and Satoh, N. (1995). Timing of Initiation of Muscle-Specific Gene-Expression in the Ascidian Embryo Precedes That of Developmental Fate Restriction in Lineage Cells. Dev Growth Differ 37, 319–327.

34. Satou, Y., Nakamura, R., Yu, D., Yoshida, R., Hamada, M., Fujie, M., Hisata, K., Takeda, H. and Satoh, N. (2019). A nearly complete genome of *Ciona intestinalis* type A (*C. robusta*) reveals the contribution of inversion to chromosomal evolution in the genus *Ciona*. Genome biology and evolution 11, 3144–3157.

35. Satou, Y., Takatori, N., Yamada, L., Mochizuki, Y., Hamaguchi, M., Ishikawa, H., Chiba, S., Imai, K., Kano, S., Murakami, S. D., et al. (2001c). Gene expression profiles in Ciona intestinalis tailbud embryos. Development 128, 2893–2904.

36. Satou, Y., Tokuoka, M., Oda-Ishii, I., Tokuhiro, S., Ishida, T., Liu, B. and Iwamura, Y. (2022). A Manually Curated Gene Model Set for an Ascidian, *Ciona robusta* (*Ciona intestinalis* Type A). Zool Sci 39.

37. Satou, Y., Yagi, K., Imai, K. S., Yamada, L., Nishida, H. and Satoh, N. (2002). macho-1-Related genes in Ciona embryos. Dev Genes Evol 212, 87–92.

38. Takahashi, H., Hotta, K., Erives, A., Di Gregorio, A., Zeller, R. W., Levine, M. and Satoh, N. (1999). Brachyury downstream notochord differentiation in the ascidian embryo. Genes Dev 13, 1519–1523.

39. Tokuoka, M., Imai, K. S., Satou, Y. and Satoh, N. (2004). Three distinct lineages of mesenchymal cells in *Ciona intestinalis* embryos demonstrated by specific gene expression. Dev Biol 274, 211–224.

40. Yagi, K., Satoh, N. and Satou, Y. (2004). Identification of downstream genes of the ascidian muscle determinant gene *Ci-macho1*. Dev Biol 274, 478–489.

41. Yagi, K., Takatori, N., Satou, Y. and Satoh, N. (2005). Ci-Tbx6b and Ci-Tbx6c are key mediators of the maternal effect gene Ci-macho1 in muscle cell differentiation in Ciona intestinalis embryos. Dev Biol 282, 535–549.

42. Yamada, A. and Nishida, H. (1999). Distinct parameters are involved in controlling the number of rounds of cell division in each tissue during ascidian embryogenesis. J Exp Zool 284, 379–391.

43. Yamada, L., Shoguchi, E., Wada, S., Kobayashi, K., Mochizuki, Y., Satou, Y. and Satoh, N. (2003). Morpholino-based gene knockdown screen of novel genes with developmental function in Ciona intestinalis. Development 130, 6485–6495.

44. Yasuo, H. and Satoh, N. (1993). Function of vertebrate T gene. Nature 364, 582–583.

45. Yasuo, H. and Satoh, N. (1994). An Ascidian Homolog of the Mouse Brachyury(T) Gene Is Expressed Exclusively in Notochord Cells at the Fate Restricted Stage. Development Growth & Differentiation 36, 9–18.

46. Yu, D., Oda-Ishii, I., Kubo, A. and Satou, Y. (2019). The regulatory pathway from genes directly activated by maternal factors to muscle structural genes in ascidian embryos. Development 146, dev173104.

